# Nitrogen fixation in mesoscale eddies of the North Pacific Subtropical Gyre: patterns and mechanisms

**DOI:** 10.1101/2021.06.03.446955

**Authors:** Mathilde Dugenne, Mary R. Gradoville, Matthew J. Church, Benedetto Barone, Samuel T. Wilson, Uri Sheyn, Matthew J. Harke, Karin M. Björkman, Nicholas J. Hawco, Annette M. Hynes, François Ribalet, David M. Karl, Edward F. Delong, Sonya T. Dyhrman, E. Virginia Armbrust, Seth John, John M. Eppley, Katie Harding, Brittany Stewart, Ana M. Cabello, Kendra A. Turk-Kubo, Mathieu Caffin, Angelicque E. White, Jonathan P. Zehr

## Abstract

Mesoscale eddies have been shown to support elevated dinitrogen (N_2_) fixation rates (NFRs) and abundances of N_2_-fixing microorganisms (diazotrophs), but the mechanisms underlying these observations are not well understood. We explored relationships among NFRs and cyanobacterial diazotroph abundances in eddy pairs of opposite polarity sampled in the North Pacific Subtropical Gyre and compared our observations to seasonal trends from the Hawaii Ocean Time-series (HOT) program. Consistent with previous reports, we found that NFRs were anomalously high for this region (up to 3.7-fold above previous monthly HOT observations) in the centers of the sampled anticyclones, coinciding with elevated abundances of *Crocosphaera* in the summertime. We then coupled our field-based observations, together with transcriptomic analyses of nutrient stress marker genes and ecological models, to evaluate potential mechanisms controlling diazotroph abundance and activity at the mesoscale. Specifically, we evaluated the role of biological (via estimates of growth and grazing rates) and physical controls on populations of *Crocosphaera*, *Trichodesmium,* and diatom symbionts. Our results suggest that increased *Crocosphaera* abundances associated with summertime anticyclones resulted from the alleviation of phosphate limitation, allowing cells to grow at rates exceeding grazing losses. In contrast, distributions of larger, buoyant taxa (*Trichodesmium* and diatom symbionts) appeared less affected by eddy-driven biological controls. Instead, they appeared driven by physical dynamics along frontal boundaries that separate cyclonic and anticyclonic eddies. Together, the interplay of eddy-specific changes in bottom-up control, top-down control, and the physical accumulation of cells likely explains the elevated diazotroph abundances and NFRs associated with anticyclones and eddy fronts.

## 2. Introduction

In the oligotrophic North Pacific Subtropical Gyre (NPSG), the supply of growth-limiting nutrients influences rates of photosynthesis in the sunlit surface ocean, determining the magnitude and variability of trophic transfer and the export of carbon (C) from the euphotic zone (Karl et al., 1996). Microorganisms with the ability to fix dinitrogen (N_2_) into bioavailable nitrogen (N), termed diazotrophs, introduce newly fixed N to otherwise N-limited phytoplankton (Karl et al., 2002). At Station ALOHA (A Long-term Oligotrophic Habitat Assessment, 22.75°N, 158.00°W), the field site of the Hawaii Ocean Time-series (HOT) program, N_2_ fixation fuels up to half of new production and N export (Böttjer et al., 2017; Karl et al., 1997). Most N_2_ fixation is controlled by unicellular diazotrophs (typically smaller than 10 *µ*m), including the cyanobacteria UCYN-A (symbionts of haptophytes), UCYN-B (*Crocosphaera*), and UCYN-C (*Cyanothece*-like organisms) (Church et al., 2009; Zehr et al., 2001). Other, larger cyanobacterial diazotrophs including filamentous *Trichodesmium* and diatom-diazotroph associations (DDAs) with the symbiotic, heterocyst-forming *Richelia* and *Calothrix* can also form episodic blooms and contribute appreciably to N_2_ fixation. Finally, non-cyanobacterial diazotrophs are also present in the NPSG (Church, Short, et al., 2005; Farnelid et al., 2011), but the contribution of these organisms to N_2_ fixation is uncertain (Moisander et al., 2017).

N_2_ fixation rates (NFRs) at Station ALOHA are highly variable, ranging from ∼0.3 to 21 nmol N L^−1^ d^−1^ in near-surface waters (Böttjer et al., 2017). Some of this variability is seasonal: NFRs generally peak during the late summer (Böttjer et al., 2017), when an annual pulse of particle export to the deep ocean has also been putatively linked to blooms of DDAs (Karl et al., 2012; Poff et al., 2021). However, several studies have also highlighted patchiness in both NFRs and diazotroph abundances over relatively small temporal (<2 days) and spatial (<30 km) scales (Gradoville et al., 2020; Robidart et al., 2014), suggesting that submesoscale and mesoscale processes may influence diazotroph distributions in the NPSG. Such physical disturbances of the upper pelagic zone are regular features of this habitat (Rii et al., 2021); for example, over a 23-year period of near-monthly sampling by the HOT program at Station ALOHA, 31% of the sampling occasions coincided with the presence of a trackable mesoscale eddy (Barone et al., 2019).

Through uplift or depression of the pycnocline, eddies perturb microbial populations by altering nutrient and light fields. As a result, mesoscale eddies, especially cyclonic eddies that drive upward displacement of isopycnal surfaces (together with associated nutrients) across the upper ocean light gradient, have been linked to elevated phytoplankton biomass and high rates of primary production in the open ocean (McGillicuddy Jr & Robinson, 1997). There is growing evidence that diazotrophic community structure and NFRs are also modified by mesoscale eddies. High NFRs have been observed in anticyclonic eddies in many ocean regions, including the South Pacific Ocean, Indian Ocean, South China Sea, and Mediterranean Sea (Holl et al., 2007; Liu et al., 2020; Löscher et al., 2016; Rahav et al., 2013). At Station ALOHA, the highest NFRs have been observed under conditions of elevated sea surface height (characteristic of anticyclonic eddies) in summer months (Böttjer et al., 2017; Church et al., 2009). Elevated abundances of *Trichodesmium*, DDAs, and *Crocosphaera* have been reported associated with anticyclonic eddies in both the North Atlantic and North Pacific Oceans (Cheung et al., 2020; Davis & McGillicuddy, 2006; Fong et al., 2008; Olson et al., 2015; Taboada et al., 2010; Wilson et al., 2017).

These studies have highlighted multiple processes that could influence N_2_ fixation in eddies of different polarities. For example, elevated NFRs and *Crocosphaera* abundances were linked to severe N-depletion in the surface waters of anticyclones, presumably driven by the depression of isopycnal surfaces (Liu et al., 2020; Wilson et al., 2017). In addition, although poorly characterized, high rates of grazing on diazotrophs and their hosts have been reported within mesoscale eddies of both polarities (Dugenne et al., 2020; Landry et al., 2008; Wilson et al., 2017). Finally, the physical movement of water, both vertically and horizontally, due to mesoscale circulation, can lead to the physical accumulation of buoyant diazotrophs (e.g. *Trichodesmium)* (Guidi et al., 2012; Olson et al., 2015). All of these mechanisms are modulated by specific life traits (e.g. buoyancy, size, symbiosis) and potential adaptations of individual taxa, suggesting that understanding diazotrophic diversity and activity within eddies is prerequisite to identifying the mechanisms driving variability in bulk NFRs. Effectively, mesoscale and sub-mesoscale features can impact both bottom-up and top-down processes controlling diazotroph diversity and activity. The relative influence of these controls is important in determining diazotroph abundance as well as the magnitude of new N delivered to the system.

In this study, we examined cyanobacterial diazotroph abundances and NFRs associated with two pairs of mesoscale eddies (cyclones and anticyclones) sampled in the NPSG and compared these data to trends from the HOT program for regional and seasonal context. We used these observations, together with environmental data, metatranscriptomes, and ecological models, to evaluate the importance of bottom-up, top-down, and physical control mechanisms in driving the mesoscale distribution and activity of diazotroph populations.

## 3. Material and Methods

### 3.1 Characterization and sampling of eddy pairs

In 2017 and 2018, two field expeditions sampled mesoscale eddies in the region north of the Hawaiian Islands. Observations took place 26 June–15 July 2017 during the Microbial Ecology of the Surface Ocean-Simons Collaboration on Ocean Processes and Ecology cruise (MESO-SCOPE, KM1709, *R/V Kilo Moana*) and 27 March–10 April 2018 during the Hawaiian Eddy Experiment (FK180310, *R/V Falkor*). Prior to each cruise, cyclonic and anticyclonic eddies were identified via satellite-derived sea level anomalies (SLA) distributed by the Copernicus Marine Service (marine.copernicus.eu/). Daily SLA maps (Fig. 1) were corrected for the multi-year trend and seasonal cycle following Barone et al. (2019). Eddies were tracked back in time to determine the stage, age, and average SLA during the sampling period (Supporting Information S1 and Table S1).

**Figure 1:**
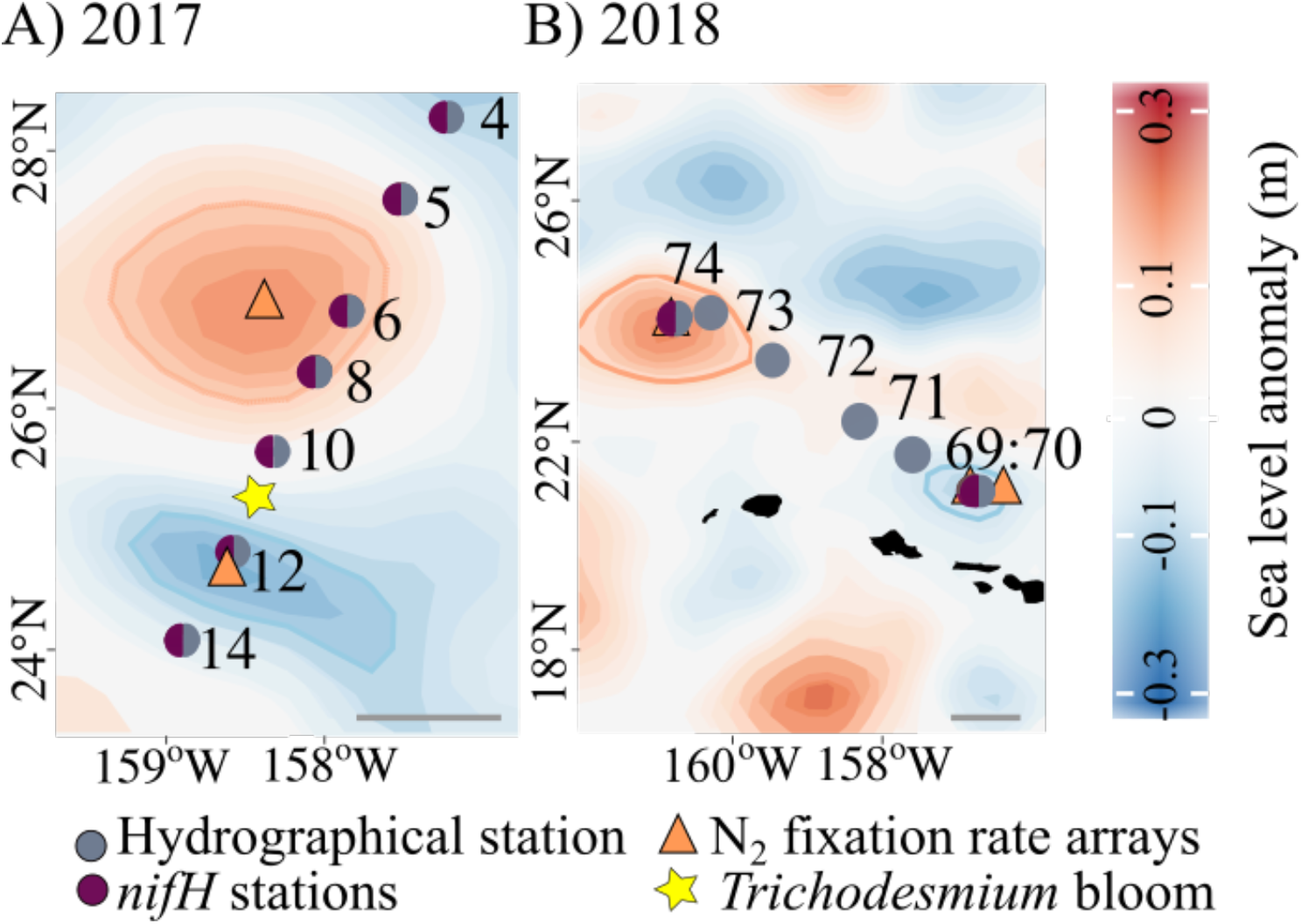
Sampling locations for *nifH* gene abundances (purple circles), hydrographical stations (blue circles), a *Trichodesmium* bloom (yellow star), and N_2_ fixation rates (triangles) across each pair of eddies during the 2017 (A) and 2018 (B) cruises. Note that surface diazotroph cell abundances were measured by autonomous flow cytometry at high spatial resolution along each cruise transect. Sampled cyclones are indicated by negative sea level anomalies (−0.1 m blue contour) and anticyclones by positive sea level anomalies (0.1 m red contour). Scale bars: 100 km.

Both cruises included an eddy mapping phase, where sampling occurred along a transect spanning one eddy to the other to characterize the water column, quantify diazotrophs via *nifH* gene abundances and automated imaging flow cytometry cell counts, and measure NFRs. They also included time-resolved, Lagrangian surveys that sampled along the drift trajectories of Surface Velocity Program (Pacific Gyre) drifters in one or both eddy centers (drogue centered at 15 m), with continuous imaging of near surface plankton communities to assess high resolution temporal dynamics.

Photosynthetically active radiation (PAR) at the sea-surface was measured using a shipboard cosine collector LI-190 (LI-COR Environmental, Lincoln, NE, USA). Downwelling PAR was measured at 1 m-intervals using a free-falling optical profiler (Satlantic HyperPro, Sea-Bird Scientific, Bellevue, WA, USA), and corrected by a factor of 1.2 to convert downwelling irradiance into scalar irradiance (Letelier et al., 2017; Wozniak et al., 2003). Both sea surface and downwelling PAR measurements were used compute the fraction of incident PAR (%) at discrete depths.

A Conductivity, Temperature, and Depth (CTD, Sea-Bird Scientific) sensor attached to a rosette sampler was used to measure depth profiles of temperature, salinity, and derived potential density. Mixed layer depth (MLD) was calculated using a 0.03 potential density offset relative to 10 m (de Boyer Montégut et al., 2004). Seawater samples collected from discrete depths using 24 × 10 L Niskin® bottles mounted to the rosette frame were used to measure soluble reactive phosphorus (hereafter phosphate, PO_4_^3-^) concentrations (precision of ± 1 nmol L^-1^ and detection limit of 3 nmol L^-1^) (Karl & Tien, 1992) and nitrate + nitrite (NO_3_^-^+NO_2_^-^) concentrations (detection limit of 1 nmol L^-1^) (Foreman et al., 2016).

Dissolved Fe (dFe) for the 2017 cruise was collected using a trace metal clean sampling rosette with external spring bottles (Ocean Test Equipment). For the 2018 cruise, samples for dFe were collected in Go-Flo® bottles mounted on a non-metallic line and triggered with a PTFE messenger. Dissolved iron concentrations were determined by inductively coupled plasma mass spectrometry (ICP-MS) using isotope dilution (Hawco et al., 2021; Pinedo-González et al., 2020). The accuracy of these measurements was confirmed by analyzing reference seawater samples, GS (0.55 ± 0.01 nM Fe, n = 3) and GSP (0.19 ± 0.03 nM Fe, n= 3), provided by the GEOTRACES community (www.geotraces.org/standards-and-reference-materials).

### 3.2 N_2_ fixation rates

NFRs were measured on all cruises using a modification of the ^15^N_2_ uptake method of Montoya et al. (1996), in which the ^15^N_2_ tracer was added as ^15^N_2_-enriched seawater to avoid incomplete dissolution of ^15^N_2_ gas (Mohr et al., 2010; Wilson et al., 2012). Prior to each cruise, surface seawater was collected from Station ALOHA, filtered through a 0.2 μm in-line filter (AcroPak 1000 capsule, Pall Corporation, Port Washington, NY, USA) and transported in the dark to the laboratory. There, the seawater was degassed, injected with ∼13 mL ^15^N_2_ gas (99 atom %, Cambridge Scientific, Watertown, MA, USA) per L seawater, and manually agitated to dissolve the gas bubble, as described by Wilson et al. (2012). The resulting ^15^N_2_-enriched seawater was dispensed into crimp-sealed, glass serum bottles, stored at 4°C for less than one week, and brought to sea. To initiate incubations, 100 mL of seawater was withdrawn from each full 4.4 L incubation bottle and replaced with 100 mL of ^15^N_2_-enriched seawater. The ^15^N/^14^N ratio of each batch of ^15^N_2_-enriched seawater was measured using membrane inlet mass spectrometry according to Ferrón et al. (2016). These values were used to calculate the initial atom % of ^15^N_2_ for each incubation. A recent study by White et al. (2020) indicates that measuring the ^15^N_2_ atom % of the enriched seawater rather than the final inoculum may lead to inaccurate initial ^15^N_2_ atom % values, which for the enriched seawater preparation method used for the eddy and HOT observations reported in this study could have led to a <20% underestimation of final calculated rates.

NFRs were measured using free-drifting, in situ arrays deployed at eddy centers (Fig. 1). Seawater was collected pre-dawn from 5, 25, 45, 75, 100, and 125 m depths using the CTD rosette Niskin® sampling bottles. Seawater was subsampled into triplicate 4.4 L acid-washed, seawater-rinsed polycarbonate bottles which were spiked with ^15^N_2_ tracer (see above), attached to the free-floating arrays, and incubated for 24 h at the depth from which they had been collected. Incubations were terminated by filtering (via a peristaltic pump or vacuum filtration) through pre-combusted (5 h at 450°C) 25 mm glass fiber filters (GF/F, Whatman, Maidstone, UK). Additional 4.4 L seawater samples were collected from each sampling depth for δ^15^N natural abundance measurements; these samples were immediately filtered at the start of each incubation. All filters were frozen at −20°C or −80°C and transported to the laboratory, where they were dried at 60°C overnight and pelleted into tin capsules (Costech Analytical Tech Inc, Valencia, CA, USA). Concentrations and isotopic composition (δ^15^N) of particulate N were analyzed via continuous-flow isotope ratio mass spectrometry using a Carlo-Erba EA NC2500 coupled with Thermo Finnigan DeltaPlus XP at the University of Hawaiʻi Biogeochemical Stable Isotope Facility. NFRs and detection limits were calculated according to Montoya et al. (1996) (Table S2).

Time-series NFRs measured at Station ALOHA are also presented to provide context for the two eddy cruises. A subset of the time-series (2012-2014) has previously been published by Böttjer et al. (2017); for this study, we extend that time series through 2019. These extended time-series measurements followed procedures identical to those described in Böttjer et al. (2017).

### 3.3 Diazotroph abundances

#### 3.3.1. DNA extraction and nifH gene quantification

The *nifH* genes of seven major cyanobacterial diazotroph groups were quantified using digital droplet polymerase chain reaction (ddPCR). During the 2018 cruise, seawater for subsequent ddPCR analyses was collected from depth profiles at the same locations sampled for ^15^N_2_ incubations. During the 2017 cruise, seawater for ddPCR was collected from depth profiles within each eddy, but from separate locations than those used for the ^15^N_2_ arrays (Fig. 1). For both cruises, seawater samples were collected from a rosette sampler and subsampled into duplicate (2018) or single (2017) 4.4 L acid-washed, milliQ-rinsed polycarbonate bottles. A peristaltic pump was used to filter 4 L of seawater through 0.2 μm pore-size Supor® membranes (Pall), which were placed into microcentrifuge tubes containing mixtures of 0.1- and 0.5-mm glass beads (Biospec Products, Bartlesville, OK, USA), flash-frozen in liquid N_2_, and stored at −80°C until analysis.

DNA was extracted from filters with a QIAcube instrument (Qiagen, Venlo, Netherlands) using the DNeasy Plant Mini Kit. Extractions followed the manufacturer’s protocol with additional steps of three flash freeze/thaw cycles, bead-beating for two min, and a Proteinase K treatment, as described by Moisander et al. (2008). The final DNA elution volume was 100 *μ*L. DNA extracts were stored at −20°C prior to ddPCR. Duplicate ddPCR reactions were performed for each sample using primers and probes targeted the following groups: *Trichodesmium* (Church, Jenkins, et al., 2005), UCYN-A (targeting the small UCYN-A1 sublineage, Church, Jenkins, et al., 2005), *Crocosphaera* (UCYN-B, Moisander et al., 2010), *Cyanothece*-like organisms, (UCYN-C, Foster et al., 2007), *Richelia* associated with *Rhizosolenia* (Het-1, Church, Short, et al., 2005), *Richelia* associated with *Hemiaulus* (Het-2, Foster et al., 2007), and *Calothrix* associated with *Chaetoceros* (Het-3, Foster et al., 2007). Descriptions of ddPCR reaction setup, droplet generation, thermocycling, thresholding, and detection limit calculations are provided in Gradoville et al. (2021).

For temporal context, we also report *nifH* gene abundances of UCYN-A, UCYN-B, *Trichodesmium*, Het-1, Het-2, and Het-3, measured at Station ALOHA between 2004 and 2017 (dataset doi: 10.5281/zenodo.4477269). For these measurements, *nifH* gene abundances were determined using quantitative PCR (qPCR); details on sample collection, DNA extraction, and qPCR are provided in Church et al. (2009). All qPCR assays except for Het-2 and Het-3 used the same primer/probe sets as the corresponding ddPCR assays for the eddy cruises. Het-2 was quantified using the primer/probes sets of Church et al. (2008), while Het-3 used the same forward and reverse primers used for ddPCR, but the probe sequence contained an extra adenine on the 3’ end.

#### 3.3.2. Diazotroph cell and filament enumeration by autonomous flow cytometry

Concentrations of large (>4 *µ*m in diameter or length, imposed by the size detection limit of the instrument) free-living and filamentous diazotrophs, including *Trichodesmium* (single filaments), *Richelia* (free and symbiotic heterocyst-containing filaments), *Calothrix* (unattached and epiphytic heterocyst-containing filaments), and large-size *Crocosphaera-like* organisms (free-living, hereafter referred as *Crocosphaera*), were estimated by automated imaging flow cytometry equipped with a 635 nm red laser (Imaging FlowCytobot [IFCb], McLane, East Falmouth, MA, USA) during the 2017 and 2018 cruises. Five mL samples were collected every 20 min from the ship’s uncontaminated underway system (∼7 m depth). Discrete bucket samples from a visual *Trichodesmium* bloom were also analyzed on the IFCb in 2017 (Fig. 1). During the analysis, individual images of single or colonial cells were collected along with their optical fingerprints: light scattering and red fluorescence (680±30 nm), emitted by chlorophyll and phycocyanin pigments. The latter is characteristic of N_2_-fixing cyanobacteria (Boatman et al., 2018; Webb et al., 2009; Zeev et al., 2008). A training set of images was used to classify organisms to the genera level based on morphological traits, as described in Dugenne et al. (2020). The output of the random forest classifier (Sosik & Olson, 2007) was corrected by manually annotating misclassified images to provide accurate estimates of cell and filament concentration of individual genera. To compare the cell counts of rare filamentous diazotrophs with *nifH* abundances, we binned consecutive samples in 2-h-intervals (in order to integrate a larger volume and lower the detection limit) and manually estimated the number of cells per filament.

Small-size *Crocosphaera* (2-4 *µ*m) were enumerated using an autonomous SeaFlow flow cytometer (Ribalet et al., 2019). Briefly, the SeaFlow was used to continuously sample surface seawater from the ship’s uncontaminated underway system, acquiring optical measurements of light scatter, red fluorescence, and orange fluorescence (characteristic of the phycoerythrin-containing *Crocosphaera*) at 3 min intervals. The population of small-size *Crocosphaera* was gated over time to estimate its abundance during time points close (less than 2 h) to collection times of the discrete *nifH* gene abundance samples.

### 3.4. Metatranscriptomic analysis of stress-marker genes

For insight into possible diazotroph growth limitation by P and Fe in eddy centers, we examined transcriptional patterns in P- and Fe-stress marker genes from samples collected during the 2017 cruise. Metatranscriptome samples were collected within the mixed layer (15 m) and the deep chlorophyll maximum (DCM, located at depths of 106±5 m and 123±5 m in the cyclone and anticyclone, respectively) at ∼4 h intervals for three days during the Lagrangian sampling of the cyclonic and anticyclonic eddies (n=18 for each eddy). Seawater samples were collected using the rosette sampler and seawater (1 L) was filtered through 25 mm diameter, 0.2 μm Supor® membranes (Pall) housed in Swinnex^TM^ units (MilliporeSigma) using a peristaltic pump. Filtration times were limited to 15-20 min. Immediately following filtration, filters were placed in RNALater (Invitrogen, Waltham, MA, USA) and stored at **-**80°C until processed.

Sample and data processing were performed using the methods described by Gifford et al. (2016) and Wilson et al. (2017). Briefly, sample processing steps included RNA extraction, spiking with internal controls, cDNA synthesis, and sequencing using Illumina NextSeq 500 v2. Approximately two million 150 bp paired-end reads were produced for each sample; raw sequences are available on NCBI SRA under project number PRJNA596510. Reads were screened for quality, trimmed, and assembled after removing sequencing primers. Genes were mapped to the ALOHA 2.0 gene catalog (Luo et al., 2020), and annotated using the Genome Taxonomy Database (GTDB, Parks et al., 2018). Raw sequence reads are available on NCBI SRA under project number PRJNA596510. Transcript counts were normalized to internal standards to account for methodological biases among samples, including library preparation and sequencing, and also normalized to the volume of water filtered for each sample as previously described (Wilson et al., 2017). Values above detection limits were further normalized to the total sample expression sum within each genus-level annotation. This step was required in order to compare eddy differential gene expression based on cellular regulation rather than based on the total number of cells. Transcripts of P- and Fe-stress marker genes (genes with higher expression levels under low P and Fe concentrations, as reviewed by Stenegren (2020) and Snow et al. (2015) were extracted from the dataset according to functional annotations assigned by comparison to the Kyoto Encyclopedia of Genes and Genomes.

Several genes exhibited diel expression patterns. To account for the expression periodicity and evaluate the differences in gene transcript abundances between eddies, we normalized the abundance of transcripts above detection limits within each eddy to the overall average of each gene transcript from the entire time-series (time points with undetected gene transcripts were excluded from this analysis). This step facilitated comparisons of different gene transcripts having large variations in their diel baseline and maximum transcript expression levels.

The statistical significance of the relative change in expression was tested using a Kruskal-Wallis test (R function ‘kruskal.test’). All gene transcripts from UCYN-A and *Calothrix*, as well as the transcripts of several genes from *Richelia* and *Trichodesmium*, were excluded from analyses as their abundances always fell below the detection limit in at least one eddy, especially in samples collected at the DCM.

### 3.5. Diazotroph populations models

#### 3.5.1. Bottom-up control on diazotroph populations

We used an ecophysiological model (Follett et al., 2018; McGillicuddy Jr, 2014; Stukel et al., 2014) to evaluate how measured abiotic factors (nutrient concentrations, light, and temperature) may have influenced diazotroph intrinsic growth rates (*µ*; per day) across the pairs of eddies. Predictions are based on the ecophysiological parameters of cyanobacterial diazotroph taxa derived from laboratory culture studies examining the effect of temperature (T), average instantaneous scalar PAR (E), PO_4_^3-^, and dFe concentrations on growth rates (Table S3). Model assumptions and caveats are described in Supporting Information S2.

We integrated the effects of abiotic factors on population growth rates by scaling the theoretical maximum growth rate of each taxon (*µ*_max_) to the relative change in growth rate (f_µ_) predicted by the main limiting factor, according to Liebig’s law of the minimum:

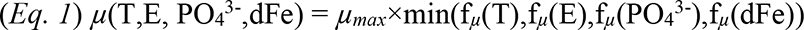

where f*_µ_* designate unitless limiting functions (Eq. 2-3, 5) representing the changes of diazotroph growth rates relative to their maximum. The change in relative growth rate as a function of T followed Eq. 2:

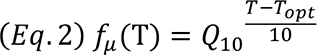

with T_opt_ (°C) the optimal temperature for which f*_µ_* (T_opt_)=1, and Q_10_, the Arrhenius coefficient corresponding to the change of growth rate over a 10°C temperature increase (reported in Table S3 for individual diazotroph taxa).

The change in relative growth rate as a function of E (*µ*mol quanta m^-2^ s^-1^) was predicted using a modification of the hyperbolic tangent function described in Jassby and Platt (1976):

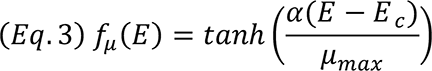

with E_c_ (*µ*mol quanta m^-2^ s^-1^), the compensation light intensity. The light saturation parameter, E_k_ (*µ*mol quanta m^-2^ s^-1^), is then derived using the maximum growth rate, *µ*_max_, reported in Table S3 and the initial slope of *µ*-E curve presented in the literature, α, as:

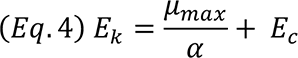

Finally, Michaelis-Menten kinetics were used to express the change in relative growth rate as a function of both PO_4_^3-^ and dFe concentration (nmol L^-1^):

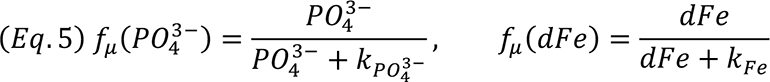

All models were fitted to published culture data using the nonlinear least-squares R function ‘nls’. We tested the differences in intrinsic growth rate estimates between eddies of opposite polarity by generalized linear regression, assuming a quasi-Binomial distribution (which is typically assumed for biological rates), weighted by the uncertainties of the estimates with the R function glm(*μ∼*Eddy, family=quasibinomial(link = ‘logit’),weights=1/ σ^2^(*μ*(T,E, PO_4_^3-^,dFe))), where the explanatory variable ‘Eddy’ corresponds to eddy polarity.

#### 3.5.2. Predation control on diazotroph populations

We explored how mesoscale eddies affected estimates of diazotroph grazing rates using a standard predator-prey model. The predation model relied on co-occurrence interactions between a given diazotroph (i.e. *Crocosphaera, Richelia, Calothrix, Trichodesmium*) and potential predators, based on abundance data from the IFCb (Dugenne et al., 2020). More specifically, we leveraged the IFCb images to (1) determine significant interactions within a generalized predator-prey model (Eq. 6) and (2) visualize ingested diazotrophic prey. We relied on the temporal dynamics of diazotrophic prey (x_i_, cells or filaments L^-1^) and putative predators (x_j_, cells L^-1^), measured in the eddy centers throughout the Lagrangian surveys (hence assuming that population dynamics are only driven by growth and loss rates), to identify significant interactions (a_ij_):

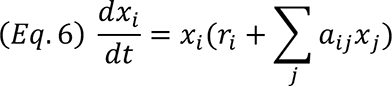

The term r comprised a linear intrinsic growth rate and a linear loss rate due to processes not explicitly accounted for with the biological interactions tested (e.g. viral lysis, programmed cell death, additional grazers not imaged by the IFCb, vertical migration, mixed layer entrainment, and/or sinking). A similar model has previously been used to assess the short-term variability of grazing rate estimates (Dugenne et al., 2020); however, here we simplified this model to have fixed parameters in each eddy center (e.g. throughout each Lagrangian survey) in order to compare grazing rates between eddies.

In the current model, the strength of the interaction between populations *i* and *j* is determined by the coefficient a_ij_ and by population abundances; a_ij_ can be interpreted as different biological rates depending on the type of the interaction. When coefficients a_ij_ and a_ji_ are negative and positive respectively, *j* grows at *i*’s expense, as would a putative grazer of the diazotroph *i*. All pairs of interaction coefficients were estimated by a linear multi-regression model fitted to the changes in population abundances throughout the Lagrangian sampling period within individual eddies as follows:

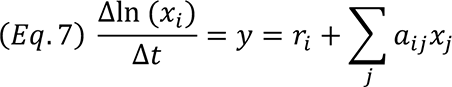

with Δt, a 1 h interval based on the sensitivity analysis presented in Supporting Information S3, yielding an average uncertainty, calculated as the ratio of the standard error over the estimate for the regression coefficient, of 12±6% of a_ij,_ the linear coefficient for the interaction with population *j*, and r_i,_ the intercept of the linear regression. Since the regression included a large number of coefficients, accommodating 103 populations in 2017 and 89 populations in 2018, we used a regularized Lasso regression implemented in the ‘glmnet’ R package to penalize non-significant interactions (Tibshirani et al., 2012). We also looked for visual evidence of diazotrophic prey ingestion using the IFCb images from 2017 and 2018 to confirm the significant interaction between diazotrophs and their putative grazers. Significant coefficients were sorted to determine the importance of individual protists based on *g_j_* (d^-1^), the product of coefficient a_ij_ and the abundance of grazer population *j*. The overall rates of grazing on a diazotroph taxon, g (d^-1^), was calculated as the sum of all the individual grazing rates:

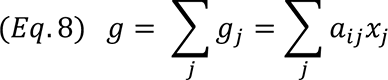

with a_ij_ negative and a_ji_ positive. The uncertainty of g was estimated by propagating errors from a_ij_ and *x_j_*.

## 4. Results

### 4.1 Mesoscale patterns in environmental conditions, NFRs, and diazotroph abundances

#### 4.1.1 Physical and biogeochemical setting

Sampled eddies varied by age and phase, as well as by in situ light, temperature, and nutrient concentrations. At the time of sampling, both anticyclones were in a stable phase (defined by the fluctuations of SLA amplitude in the eddy centers, Table S1) and both cyclones were in a weakening phase. The two pairs of eddies were sampled in different seasons: summer (July) in 2017 and early spring (March-April) in 2018. Physical variables generally followed expected seasonal trends, with higher sea surface temperature and daily incident light in the 2017 cruise (Fig. 2) and the deepest mixed layer observed in the 2018 cruise (2018 anticyclone MLD = 50.3 ± 17.8 m; mean MLD in other eddies ranged 17.8-30.6 m, Table S1). Temperature and in situ light flux decreased with depth, except in the 2018 anticyclone, where temperature was nearly constant within the upper 150 m (Fig. 2). Near-surface temperatures were similar between eddies within each cruise; however, temperatures were lower in the lower euphotic zone of the cyclonic eddies (Fig. 2), consistent with the uplift of deeper isopycnals. Daily integrated light did not vary between eddies during the 2017 cruise, while in the 2018 cruise, lower daily-integrated light was observed in the anticyclone due to cloud cover on the day of sampling (Fig. 2).

**Figure 2:**
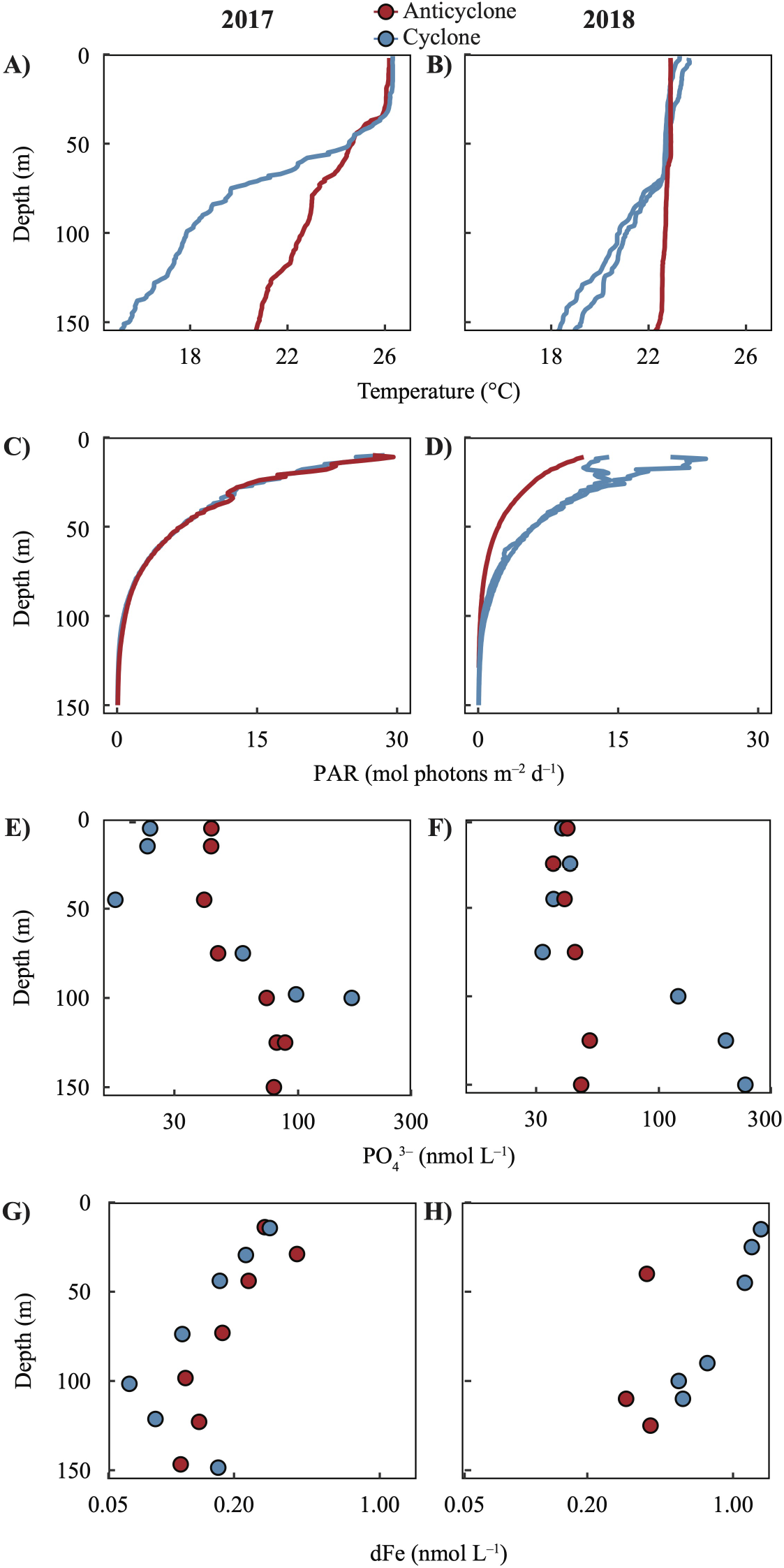
Depth profiles of temperature (A-B, linear scale), PAR (C-D, linear scale), PO_4_^3–^ concentrations (E-F, log scale), and dFe concentrations (G-H, log scale) on the 2017 (left) and 2018 (right) cruises. Red and blue indicate observations from the anticyclonic and cyclonic eddies, respectively. Profiles of temperature and PAR are from the days and locations of the N_2_ fixation measurements (note that N_2_ fixation rates were measured twice within the 2018 cyclone); PO_4_^3-^ and dFe concentrations were measured at nearby stations within the same eddy.

PO_4_^3-^ concentrations varied with depth and eddy polarity. PO_4_^3-^ concentrations were lowest at the surface and increased with depth, except for the 2018 anticyclone, in which concentrations were relatively stable within the upper 150 m, consistent with the near-homogenous temperature profile (Fig. 2) and deep downwelling intensified by winter mixing (Table S1). At depth (100-150 m), PO_4_^3-^ concentrations were always higher in cyclones than in anticyclones (Fig. 2) due to eddy-induced changes in the depths of isopycnal surfaces. In contrast, during the 2017 cruise, PO_4_^3-^ concentrations in the upper 45 m were greater in the anticyclone (42 ± 2 nmol L^−1^) than in the cyclone (21 ± 4 nmol L^−1^) (Fig. 2). Consequently, the surface [NO_3_^-^ + NO_2_^-^]:[PO_4_^3-^] ratio was lower in anticyclone (0.12 ± 0.03 mol:mol at 15 m, n=3) than in the cyclone (0.37 ± 0.4 mol:mol at 15 m, n=2) during 2017; all surface ratios were well below Redfield stoichiometry (N:P <1). During the 2018 cruise, PO_4_^3-^ concentrations (range: 35-42 nmol L^−1^) and inorganic [NO_3_^-^ + NO_2_^-^]:[PO_4_^3-^] ratios (0.15-0.34 mol:mol) in the upper 45 m did not differ between the cyclonic and anticyclonic eddies (Fig. 2).

Dissolved Fe concentrations ranged from 0.06 to 1.36 nmol L^−1^ within the upper 150 m across all eddies (Fig. 2). Concentrations were typically highest at the surface and decreased with depth into the lower euphotic zone; in the 2017 cyclone, dFe concentrations reached a minimum near the deep chlorophyll maximum and increased in deeper waters alongside the increase in macronutrients (Hawco et al., 2021). In the 2018 anticyclone, dFe concentrations did not vary within the upper 150 m, consistent with the deeper mixed layer and downwelling of surface waters in this eddy. Concentrations were generally higher in the 2018 cruise than in the 2017 cruise, and near-surface dFe patterns between eddies also varied on the two cruises. During the 2017 cruise, dFe in the upper 45 m ranged from 0.17 to 0.31 nmol L^−1^ and were slightly higher in the anticyclone (0.30 ± 0.08 nmol L^−1^) than in the cyclone (0.23 ± 0.06 nmol L^−1^). During the 2018 cruise, elevated dFe concentrations were observed in the mixed layer of the cyclone (1.24 ± 0.11 nmol L^−1^ in the upper 45 m) but not in the anticyclone (0.39 nmol L^−1^ at 40 m, within the mixed layer). Concentrations greater than 1 nmol L^−1^ in the 2018 cyclone are anomalously high compared to the existing measurements for Station ALOHA (typically between 0.2–0.5 nmol L^−1^ in the mixed layer (Fitzsimmons et al., 2015)). In the weeks prior to sampling, the path of 2018 cyclonic eddy deflected off the northern coast of Maui, possibly leading to input of dFe from coastal sources along the Hawaiian Islands.

#### 4.1.2. N_2_ fixation rates and diazotroph abundances

We measured NFRs using *in situ* arrays deployed for 24 h in each eddy center. The highest NFRs were observed in the anticyclones during both cruises, with the highest rates occurring in the 2017 anticyclone (up to 18.6 nmol N L^−1^ d^−1^, Fig. 3). There were strong differences in NFRs between eddies sampled during the 2017 cruise: depth-integrated rates (0-125 m) were ∼6 times higher in the anticyclonic eddy (670 *µ*mol N m^-2^ d^-1^) than in the cyclonic eddy (115 *µ*mol N m^-2^ d^-1^). During the 2018 cruise, depth-integrated NFRs were ∼1.7-fold higher in the anticyclone (428 *µ*mol N m^-2^ d^-1^) than the cyclone (240-267 *µ*mol N m^-2^ d^-1^) (Fig. 2). In all four sampled eddies, NFRs were highest at the surface and decreased with depth.

**Figure 3:**
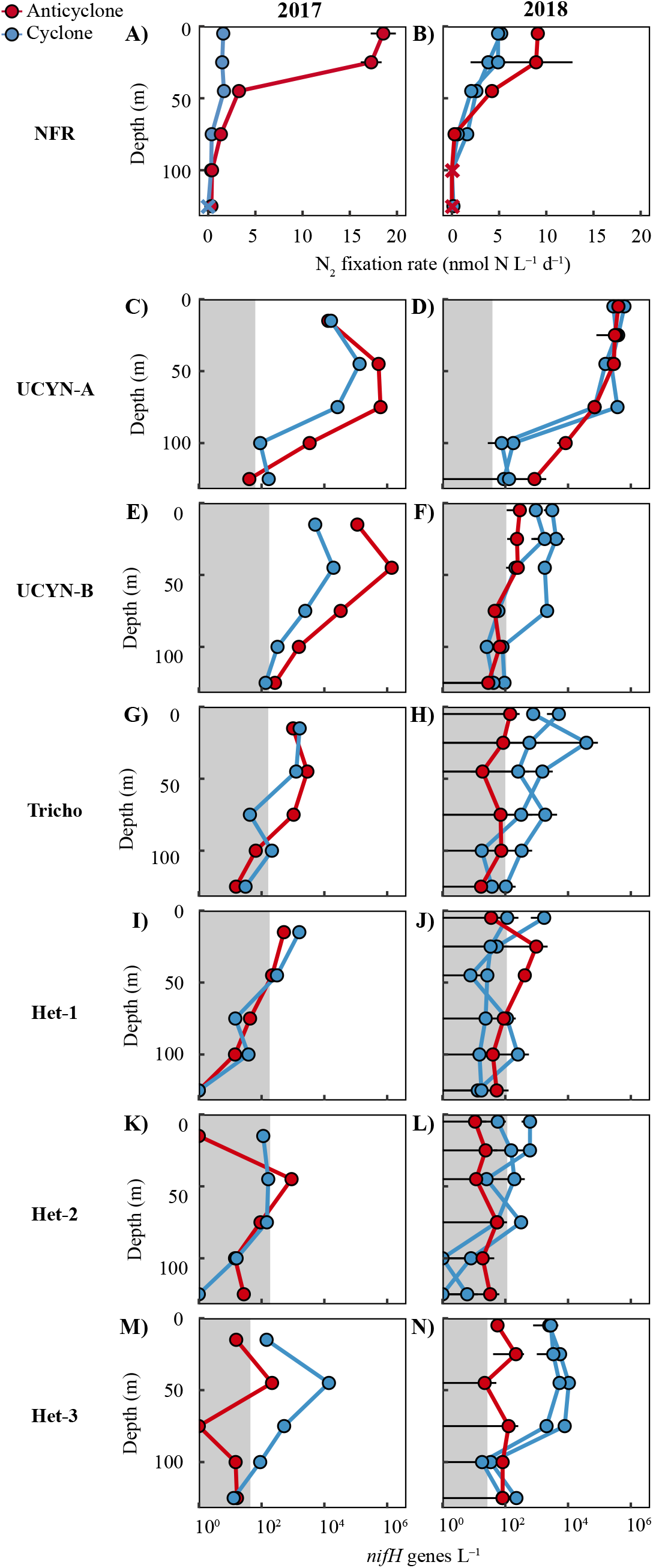
N_2_ fixation rates (NFRs, A-B, linear scale) and *nifH* gene abundances (C-N, log-scale), measured in the center of eddies during the 2017 (left) and 2018 (right) cruises. Note that NFRs and *nifH* gene abundances were measured twice within the 2018 cyclone. × represent rates below detection limits; shaded areas indicate the detection limits of each ddPCR assay. Error bars represent the standard deviation among biological replicates (n=3 for NFR data, n=2 for 2018 ddPCR data, not available for 2017 ddPCR data). *nifH* gene abundances of UCYN-C were relatively low (maximum 2.2 × 10^4^L^−1^) and are not presented here; they can be found in Supporting Information Table S4.

We estimated diazotroph abundances by quantifying the *nifH* genes of seven cyanobacterial taxa and by enumerating *Trichodesmium*, *Calothrix*, *Richelia*, and *Crocosphaera* cells/filaments using automated flow cytometry. Depth profiles of *nifH* gene abundances were measured in eddy centers (Fig. 3, Table 1, Table S4) and along a high-resolution sampling transect spanning the two eddies during the 2017 cruise (Fig. 4). Automated flow cytometry cell counts were measured continuously in the surface layer and four consecutive samples (∼2 h sampling time) corresponding to the nearest *nifH* samples were binned to provide estimates of cell abundances in eddy centers (Table 1). Together, the *nifH* gene- and cell-based datasets indicate that the small diazotrophs UCYN-A and UCYN-B (*Crocosphaera*) were numerically dominant over large diazotroph taxa in all sampled eddies.

**Figure 4:**
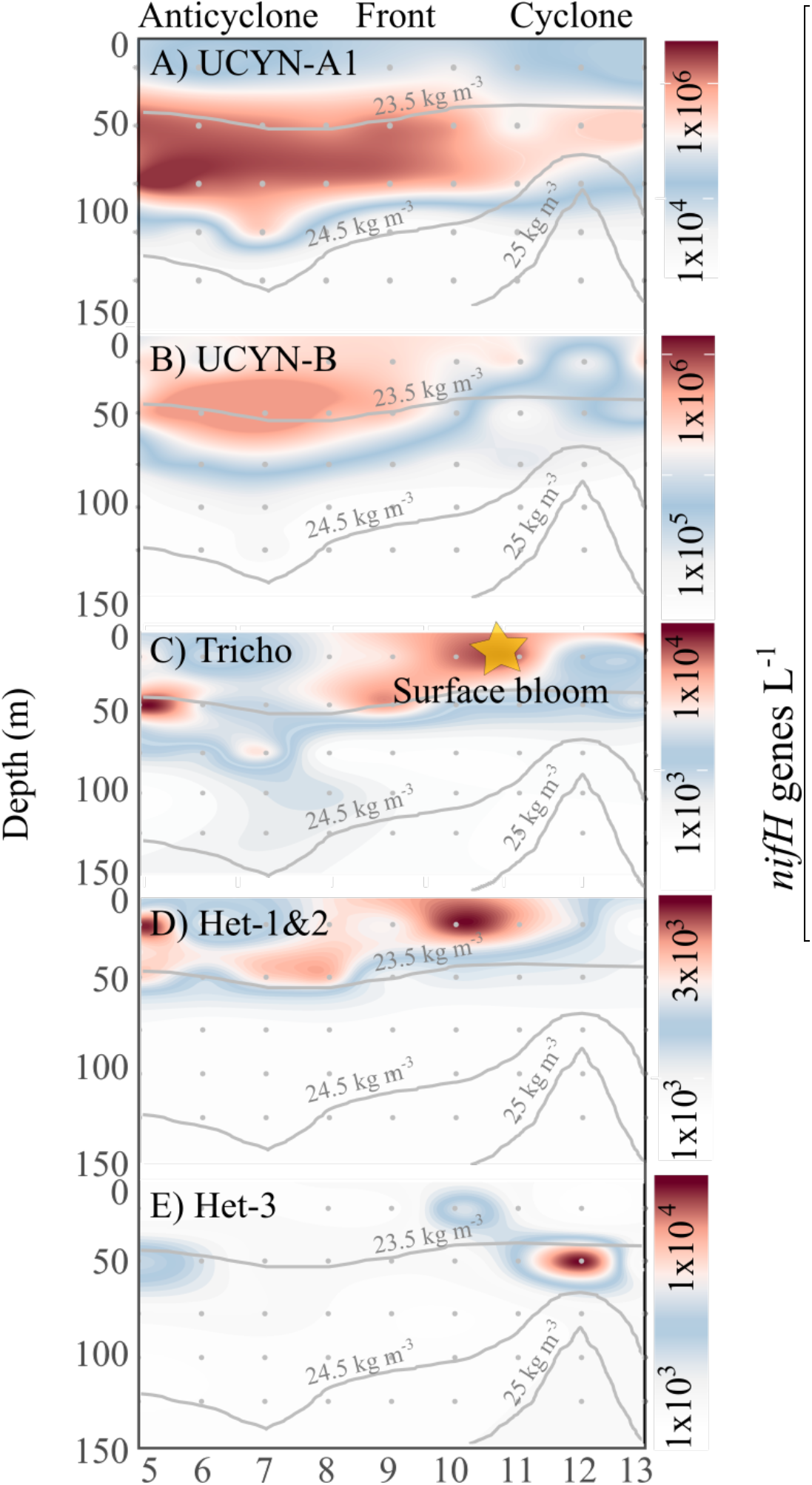
Interpolated *nifH* gene abundances of UCYN-A1 (A), UCYN-B (*Crocosphaera*, B), Tricho (*Trichodesmium*, C), Het-1&2 (*Richelia*, D), and Het-3 (*Calothrix*, E) across the high resolution 2017 eddy transect. The contours of potential density anomaly (kg m^-3^) in A-E depict the vertical displacement of isopycnal surfaces associated with mesoscale eddies. A surface bloom of *Trichodesmium* (yellow star, positioned in the contour based on its latitude between Stations 10 and 11) was visually detected and sampled with imaging flow cytometry.

**Table 1:**
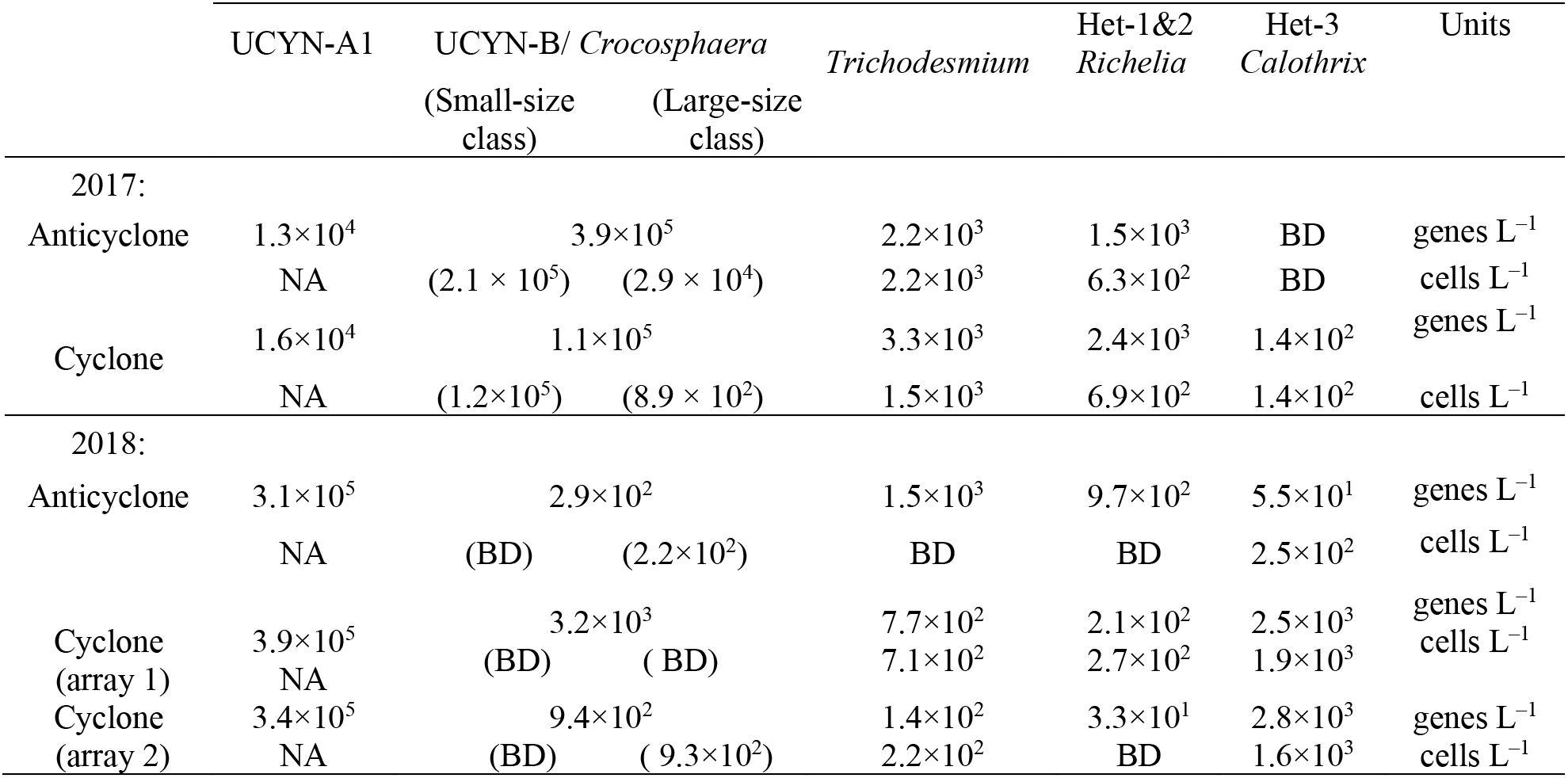
Average *nifH* gene (genes L^-1^) and cell abundances (cells L^-1^) of diazotroph taxa measured by ddPCR (15 m for 2017 cruise; 25 m for 2018 cruise) and automated flow cytometry (7 m) in eddy centers. NA: data not available. BD: below detection limits.

|  | UCYN-A1 | UCYN-B/ *Crocospaera* |  | *Trichodesmium* | Het-1&2<br>*Richelia* | Het-3<br>*Calothrix* | Units |
| --- | --- | --- | --- | --- | --- | --- | --- |
|  |  | (Small-size<br>class) | (Large-size<br>class) |  |  |  |  |
| 2017: |  |  |  |  |  |  |  |
| Anticyclone | 1.3×10<sup>4</sup> | 3.9×10<sup>5</sup> |  | 2.2×10<sup>3</sup> | 1.5×10<sup>3</sup> | BD | genes L<sup>-1</sup> |
|  | NA | (2.1 × 10<sup>5</sup>) | (2.9 × 10<sup>4</sup>) | 2.2×10<sup>3</sup> | 6.3×10<sup>2</sup> | BD | cells L<sup>-1</sup> |
| Cyclone | 1.6×10<sup>4</sup> | 1.1×10<sup>5</sup> |  | 3.3×10<sup>3</sup> | 2.4×10<sup>3</sup> | 1.4×10<sup>2</sup> | genes L<sup>-1</sup> |
|  | NA | (1.2×10<sup>5</sup>) | (8.9 × 10<sup>2</sup>) | 1.5×10<sup>3</sup> | 6.9×10<sup>2</sup> | 1.4×10<sup>2</sup> | cells L<sup>-1</sup> |
| 2018: |  |  |  |  |  |  |  |
| Anticyclone | 3.1×10<sup>5</sup> | 2.9×10<sup>2</sup> |  | 1.5×10<sup>3</sup> | 9.7×10<sup>2</sup> | 5.5×10<sup>1</sup> | genes L<sup>-1</sup> |
|  | NA | (BD) | (2.2×10<sup>2</sup>) | BD | BD | 2.5×10<sup>2</sup> | cells L<sup>-1</sup> |
| Cyclone<br>(array 1) | 3.9×10<sup>5</sup> | 3.2×10<sup>3</sup> |  | 7.7×10<sup>2</sup> | 2.1×10<sup>2</sup> | 2.5×10<sup>3</sup> | genes L<sup>-1</sup> |
|  | NA | (BD) | (BD) | 7.1×10<sup>2</sup> | 2.7×10<sup>2</sup> | 1.9×10<sup>3</sup> | cells L<sup>-1</sup> |
| Cyclone<br>(array 2) | 3.4×10<sup>5</sup> | 9.4×10<sup>2</sup> |  | 1.4×10<sup>2</sup> | 3.3×10<sup>1</sup> | 2.8×10<sup>3</sup> | genes L<sup>-1</sup> |
|  | NA | (BD) | (9.3×10<sup>2</sup>) | 2.2×10<sup>2</sup> | BD | 1.6×10<sup>3</sup> | cells L<sup>-1</sup> |

During the 2017 cruise, *nifH* gene abundances of UCYN-A (up to 1.4×10^6^ *nifH* genes L^-1^) and UCYN-B (up to 6.2×10^5^ *nifH* genes L^−1^) were higher in the anticyclone center than in the cyclone center (Table 1, Fig. 3). Independent measurements of *Crocosphaera* cells via flow cytometry likewise showed higher average concentrations in the anticyclone (2.4×10^5^ cells L^−1^) than in the cyclone (1.2×10^5^ cells L^−1^) and included the presence of both small and large sub-populations (Table 1). Abundances of other taxa were lower (maximum 5.1×10^3^ *nifH* genes L^−1^ and 2.2×10^3^ cells L^−1^ in surface waters of eddy centers, Table 1) and did not differ between eddies, except for Het-3, which had higher abundances in the cyclone than in the anticyclone. Abundances of most groups decreased with depth, although UCYN-A, UCYN-B, UCYN-C, Het-2, and Het-3 all displayed subsurface maxima (Fig. 3, Table S4).

On the transect spanning the two 2017 eddies, large and small diazotroph taxa displayed different spatial patterns relative to isopycnal displacements (Fig. 4). Abundances of *Trichodesmium* and Het-1&2 (symbionts of diatoms) were maximal in surface waters on the northern edge of the anticyclone and in surface waters on the cyclonic side of the front separating the two eddies, where a visual bloom of *Trichodesmium* was also observed (Fig. 4). Bucket samples were analyzed with the IFCb to measure the concentration of filaments at the bloom location (148 ± 44 filaments mL^−1^). Het-3 abundances were highest in subsurface (45 m) waters of the cyclonic eddy, at the base of the isopycnal uplift. In contrast, abundances of UCYN-B and UCYN-A were highest in the anticyclonic eddy, with maximal abundances from 15-45 m for UCYN-B and 45-75 m for UCYN-A (Fig. 4).

During the 2018 cruise, UCYN-A had the highest *nifH* gene abundances of all groups quantified (maximum abundance of 6.2 × 10^5^ *nifH* genes L^−1^, Table 1). UCYN-A abundances within the upper 75 m did not vary between eddy centers, while in the lower euphotic zone (100-125 m), abundances decreased but were higher in the anticyclone (4.6 × 10^3^ ± 5.6 × 10^3^ *nifH* genes L^−1^) than in the cyclone (1.2 × 10^2^ ± 4.7 × 10^1^ *nifH* genes L^−1^) (Fig. 3).

Abundances of other groups were lower (maximum 3.8×10^4^ *nifH* genes L^−1^ and 1.9×10^3^ cells L^−1^ in surface waters of eddy centers, Table 1) and generally decreased with depth (Fig. 3). *nifH* abundances of Het-1 did not display consistent differences between eddies (Fig. 3, Table S4). In comparison, UCYN-B, UCYN-C, *Trichodesmium*, Het-2, and Het-3 were all higher in the cyclonic eddy than the anticyclonic eddy, despite higher NFRs in the latter. IFCb cell count data from 2018 likewise show that abundances of large diazotrophs were generally lower than in 2017, and more abundant in the cyclone. Notably, *Crocosphaera* cells all belonged exclusively to the large-size phenotype and *Calothrix* was mainly found free-living and not attached to their diatom hosts.

### 4.2 Growth and grazing of diazotrophs at the mesoscale

We investigated potential bottom-up and top-down controls on diazotroph populations to understand the mechanisms driving observed differences in abundances and NFRs between eddies. Bottom-up control mechanisms were explored by modeling diazotroph growth rates as a function of limiting factors (Section 4.2.1) and examining patterns in diazotroph P- and Fe-stress marker gene expression in eddies (Section 4.2.2). Top-down controls were explored by comparing the community structure of putative diazotroph grazers and modeled grazing rates in the sampled eddies (Section 4.2.3).

#### 4.2.1. Modeled diazotroph growth rates and limiting factors

Diazotroph growth rates and the factors limiting growth were predicted for the euphotic zone along eddy transects. This was accomplished using a physiological model forced with mean *in situ* PAR, temperature, PO_4_^3-^, and dFe concentrations and parameterized using taxa-specific ecophysiological parameters gleaned from culture studies in the existing literature (Table S3). Due to their dependence on light and/or temperature, growth rate predictions were generally higher within the mixed layer and decreased with depth, with the exception of DDAs, for which the effect of temperature has not been tested in cultures (Supporting Information S2). Growth rates of UCYN-A could not be modeled due to a lack of available culture data and growth parameterization.

In general, the compilation of growth parameters highlighted large differences in theoretical maximum growth rates among taxa, with values of 0.86 d^-1^, 0.6 d^-1^, 0.26 d^-1^, and 0.33 d^-1^ for large-size *Crocosphaera*, *Richelia*, *Calothrix*, and *Trichodesmium*, respectively. Growth rate predictions across eddy transects were always highest for *Crocosphaera*, with maximum rates of 0.81 ± 0.20 d^-1^ and 0.43 ± 0.07 d^-1^ within the mixed layer in 2017 and 2018 respectively (Fig. 5A, G, Table 2). *Trichodesmium* appeared to grow at rates that were near its theoretical maximum in 2017 (0.28-0.29 d^-1^) and slower rates in 2018 presumably due to lower observed temperatures (Fig. 2, Fig. 5K, L). In comparison, predictions for DDAs were consistently lower than their theoretical maximum, with maximum growth rates of 0.25 d^-1^ and 0.13 d^-1^ for *Richelia* and *Calothrix* respectively. DDA growth rates were generally higher in 2018, when nutrient concentrations were elevated, and at depth (∼50-75 m) in 2017, when subsurface nutrient concentrations were well above that of the stratified surface layer (Fig. 2).

**Figure 5:**
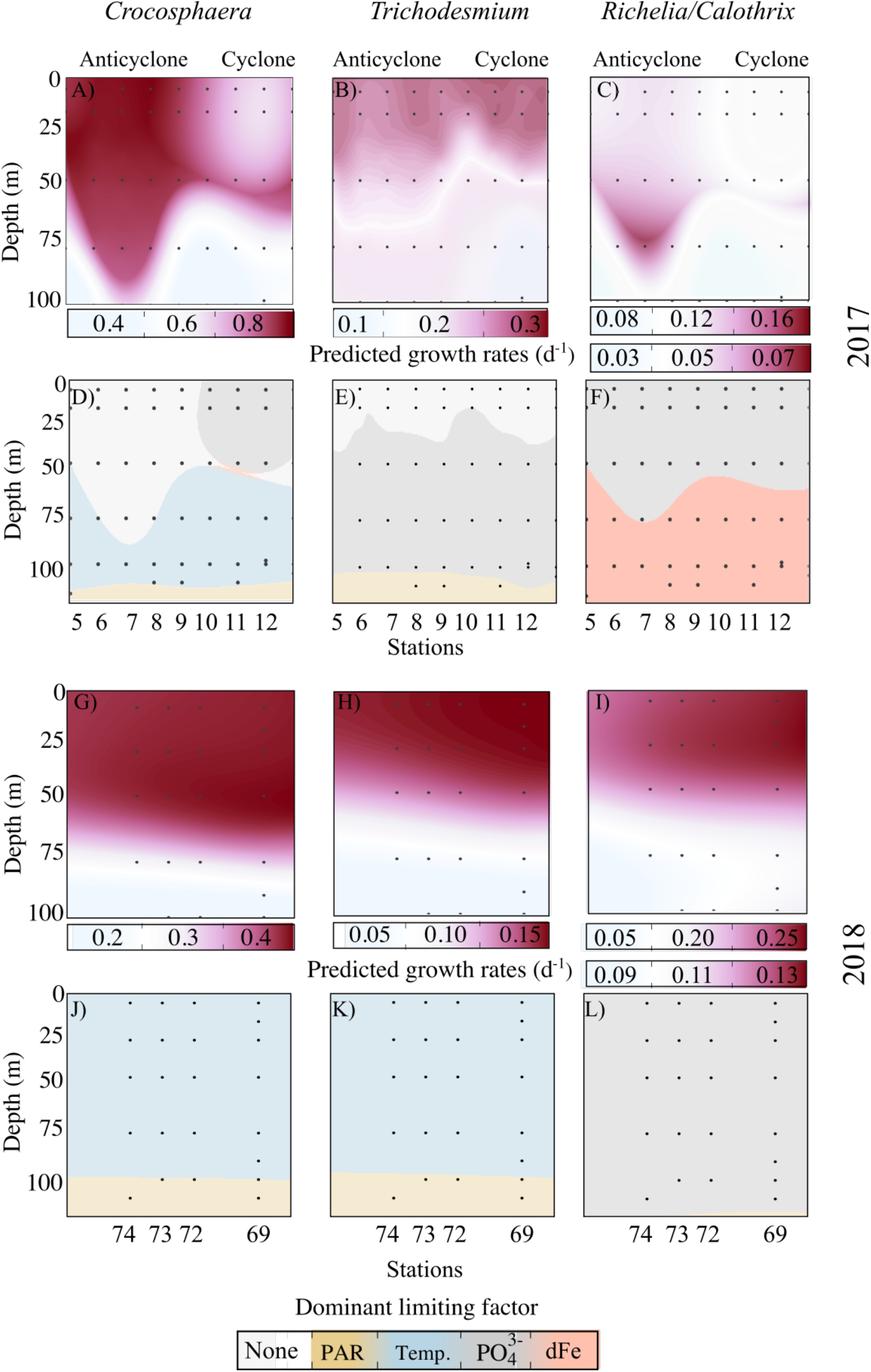
Estimates of the growth rates and limiting factors for *Crocosphaera* (A, D, G, J), *Trichodesmium* (B, E, H, K), and *Richelia/Calothrix* (C, F, I, L) in eddy cruises. Predictions of growth rates (A-C, G-I) and limiting factors (D-F, J-L) across the eddy transect are based on parameters provided in Table S3 combined with interpolated cross-sections of temperature (Temp.), average scalar irradiance (PAR), and inorganic nutrient concentrations (PO_4_^3-^, dFe). Since parameters for limiting resources were identical for *Richelia* and *Calothrix*, with the exception of respective maximum growth rates reflected in the top (*Richelia*) and bottom (*Calothrix*) scales, predictions showed similar patterns. The locations of the observations with the lowest resolution (nutrient concentrations) are indicated by the black dots.

**Table 2:**
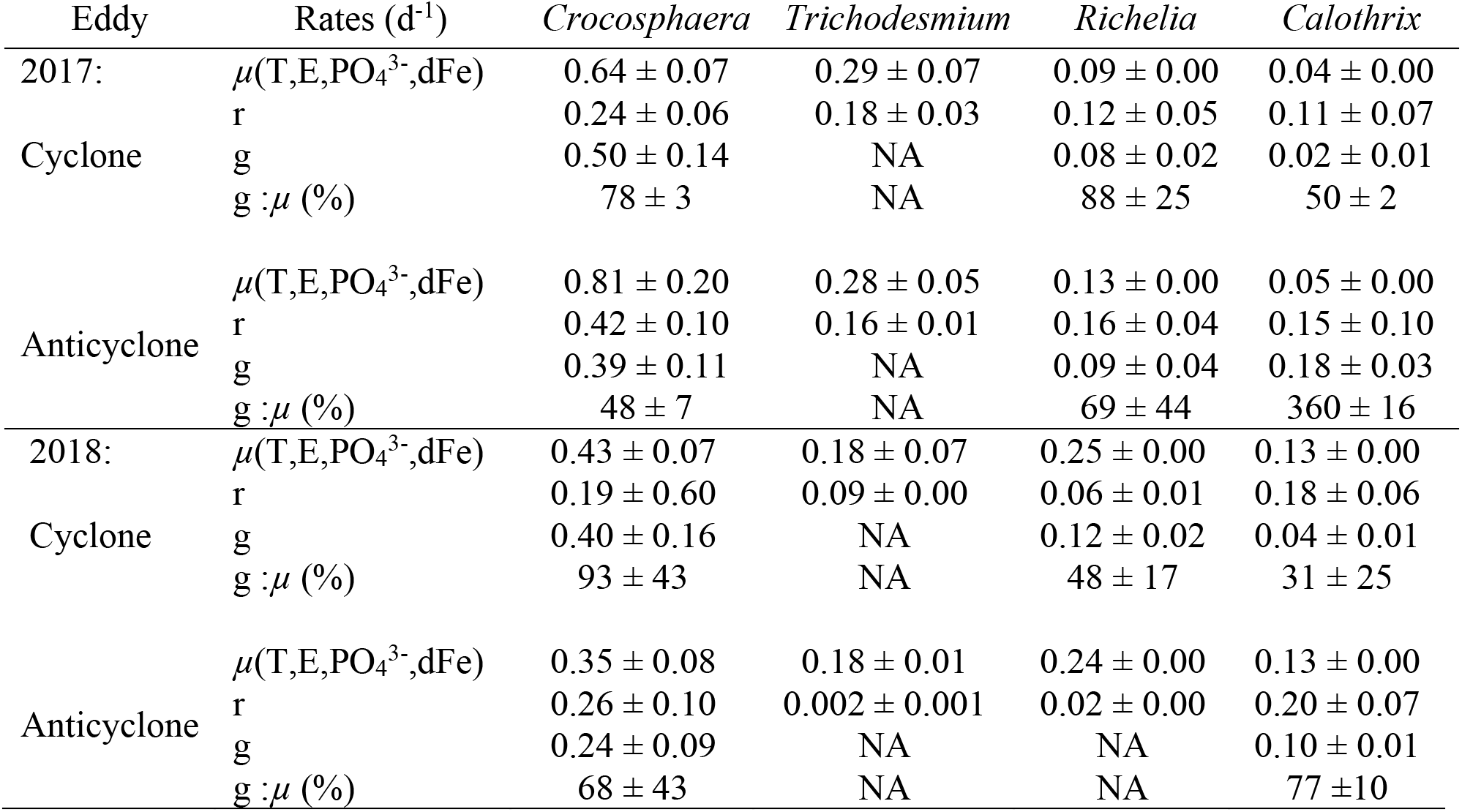
Predictions of diazotroph biological rates in eddy centers during the 2017 and 2018 cruises. Biological rates were derived from population models applied to diazotrophs imaged by the IFCb (large *Crocosphaera*, *Trichodesmium*, *Richelia*, and *Calothrix*). Intrinsic growth rates, *µ*(T,E, PO_4_^3-^,dFe), were predicted across the euphotic zone based on culture adaptations to temperature (T), instantaneous light (E), phosphate (PO_4_^3-^) concentration, and dissolved iron (dFe) concentration (Fig. 5). Here, we report growth rates predictions (*µ*(T,E, PO_4_^3-^,dFe) ± standard deviation derived from the uncertainty of parameters) matching estimates of surface (7m) diazotroph growth rates (r ± standard deviation) and grazing rates (g ± standard deviation) based on Lagrangian time-series of diazotroph and protistan populations. The grazing pressure (g:*µ*) is reported to indicate whether a diazotroph population grew faster than it was grazed upon (g<*µ*), allowing for potential accumulation. Note that r is generally lower than the intrinsic growth rates *µ*(T,E, PO_4_^3-^,dFe), as this parameter accounts for additional losses from other grazers, viruses, programmed cell death, vertical migration, and/or sinking. NA: Not Available

Predicted dominant limiting factors varied among taxa and between sampling periods (Fig. 5D-F, J-L). Given the relatively low affinity of *Crocosphaera* for PO_4_^3-^ (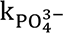=75 ± 12 nmol L^-1^) and the large differences in surface concentrations observed in 2017 (Fig. 2), predicted surface growth rates (≤50 m) appeared PO_4_^3-^-limited in the cyclone and not limited by any factor in the anticyclone (based on a limitation threshold of 80% of the maximum theoretical growth rate). Consequently, predicted *Crocosphaera* growth rates were significantly higher in the 2017 anticyclone (0.81 ± 0.20 d^-1^) than in the 2017 cyclone (0.64 ± 0.07 d^-1^). This represents the only significant difference in predicted surface growth rates between eddies across all taxa examined. However, predicted dominant limiting factors did vary by taxa and year. In 2017, *Trichodesmium* appeared non-limited at the surface and equally limited by PO_4_^3-^ between 50-100 m and by PAR below 100 m, while *Richelia/Calothrix* appeared PO_4_^3-^-limited at the surface (≤50 m) and dFe-limited at depth (≥75 m) in both eddy centers. In 2018, the dominant limiting factors for *Crocosphaera* and *Trichodesmium* were predicted to be temperature above 100 m and PAR at deeper depths, while DDAs were predicted to be PO_4_^3-^-limited throughout the water column. None of the surface or deep growth rate estimates differed between eddies, as all abiotic factors examined remained similar in the cyclone and anticyclone (Fig 2).

#### 4.2.2. Diazotroph expression of P- and Fe-stress marker genes

We analyzed patterns of diazotroph expression of known P- and Fe-stress marker genes using metatranscriptomes collected within the mixed layer and deep chlorophyll maximum during the 2017 cruise. Transcripts of six P-stress genes and three Fe-stress genes were detected for *Crocosphaera* in surface waters (Fig. 6). Fewer gene transcripts were detected for *Trichodesmium* and *Richelia* in surface waters, and transcript abundance for all these diazotrophic cyanobacterial groups were minimal at the DCM, as expected given their lower abundances (Fig. 6). No genes examined in UCYN-A or *Calothrix* had transcript levels above detection limits. *Crocosphaera* transcript abundances of all six P-stress marker genes were significantly higher within the mixed layer of the cyclonic eddy, where PO_4_^3-^ concentrations were significantly lower (Fig. 2), as compared to the anticyclonic eddy (Fig. 6). Transcripts for two *Richelia* P-stress genes were detected at the surface: the sulfolipid biosynthase (*sqdB)* abundances were significantly higher in the cyclonic eddy, while the high-affinity phosphate transporter (*pstS)* abundances did not differ between eddies. The only *Trichodesmium* P-stress gene transcript detected, a phosphonate transporter (*phnE)*, did not differ in abundance between eddies.

**Figure 6:**
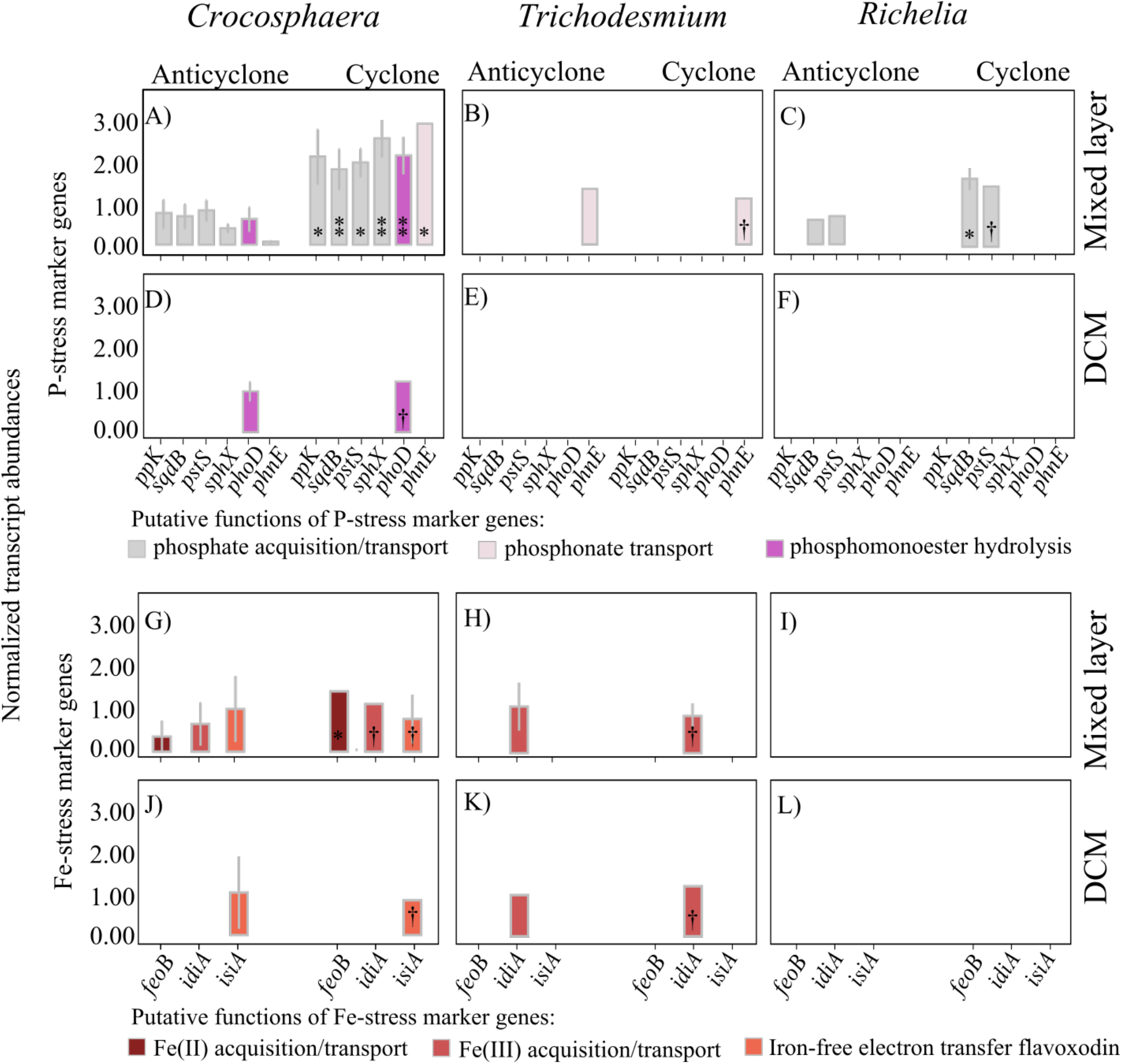
Relative expression of P-(A-F) and Fe-stress (G-L) marker genes (according to Stenegren (2020) and Snow et al. (2015)) of *Crocosphaera*, *Trichodesmium*, and *Richelia* within the mixed layer (15 m) and at the deep chlorophyll maximum (DCM) in 2017. Individual bars represent the average (± standard errors) standardized transcript abundances for each stress marker gene in a specific eddy. P-stress markers include genes related to dissolved inorganic (*ppk: polyphosphate kinase, sqdB: sulfolipid biosynthase (substitution of P-lipids), pstS/sphX: high-affinity phosphate transporter*) and organic (*phoD: Alkaline phosphatase, phnE: phosphonate transporter*) P acquisition/transport. Fe-stress markers include genes related to dissolved inorganic iron acquisition/transport (*feoB: ferrous iron transporter, idiA: iron deficiency-induced protein*) and the substitution of enzymes involved in electron transport (*isiA: flavodoxin*)*. P*-values: ** (<0.005) * (<0.01) † (not significant)

Of the three *Crocosphaera* Fe-stress genes with transcripts detected in the mixed layer, only one (*feoB*, related to Fe(II) acquisition/transport) had higher transcripts levels in the cyclonic eddy (Fig. 6G). The other two detected gene transcripts (*idiA*, the putative periplasmic Fe(III) binding protein, and *isiA*, encoding the iron-free electron transfer protein flavodoxin) had no significant difference in transcript abundance between eddies (Fig. 6G, J). The only additional diazotroph Fe-stress gene transcript detected was the iron deficiency-induced protein (*idiA)* gene for *Trichodesmium*, which had no significant difference in abundance between eddies (Fig. 6H,K).

#### 4.2.3. Diazotroph grazer community structure and estimated grazing rates

We tracked the abundances of several putative grazers of cyanobacterial diazotrophs using the IFCb. Diazotroph grazers were identified by analyzing protists co-occurring with *Crocosphaera*, *Trichodesmium*, or DDAs, and via microscopic visual evidence of diazotroph ingestion in IFCb images (Fig. 7). Putative grazers included the dinoflagellates *Gymnodinium, Cochlodinium*, *Protoperidinium, Katodinium,* and *Prorocentrum*, the ciliates *Strombidium* and *Uronema,* as well as large copepods and nauplii, most of which were imaged with ingested diazotrophs during the eddy cruises (Fig. 7A-I). Results of a predator-prey model provide estimates of grazing rates on specific diazotrophs during the two eddy cruises (Fig. 7J-K). The grazing rates of copepods were not accounted for in this analysis, as their low abundance precluded the predator-prey model from accurately estimating grazing rates (Supporting Information S3). Also, grazing by small protists (<4 *μ*m) or upon small diazotrophs (<4 *μ*m, including UCYN-A) could not be assessed using this method due to the size constraints of IFCb imaging.

**Figure 7:**
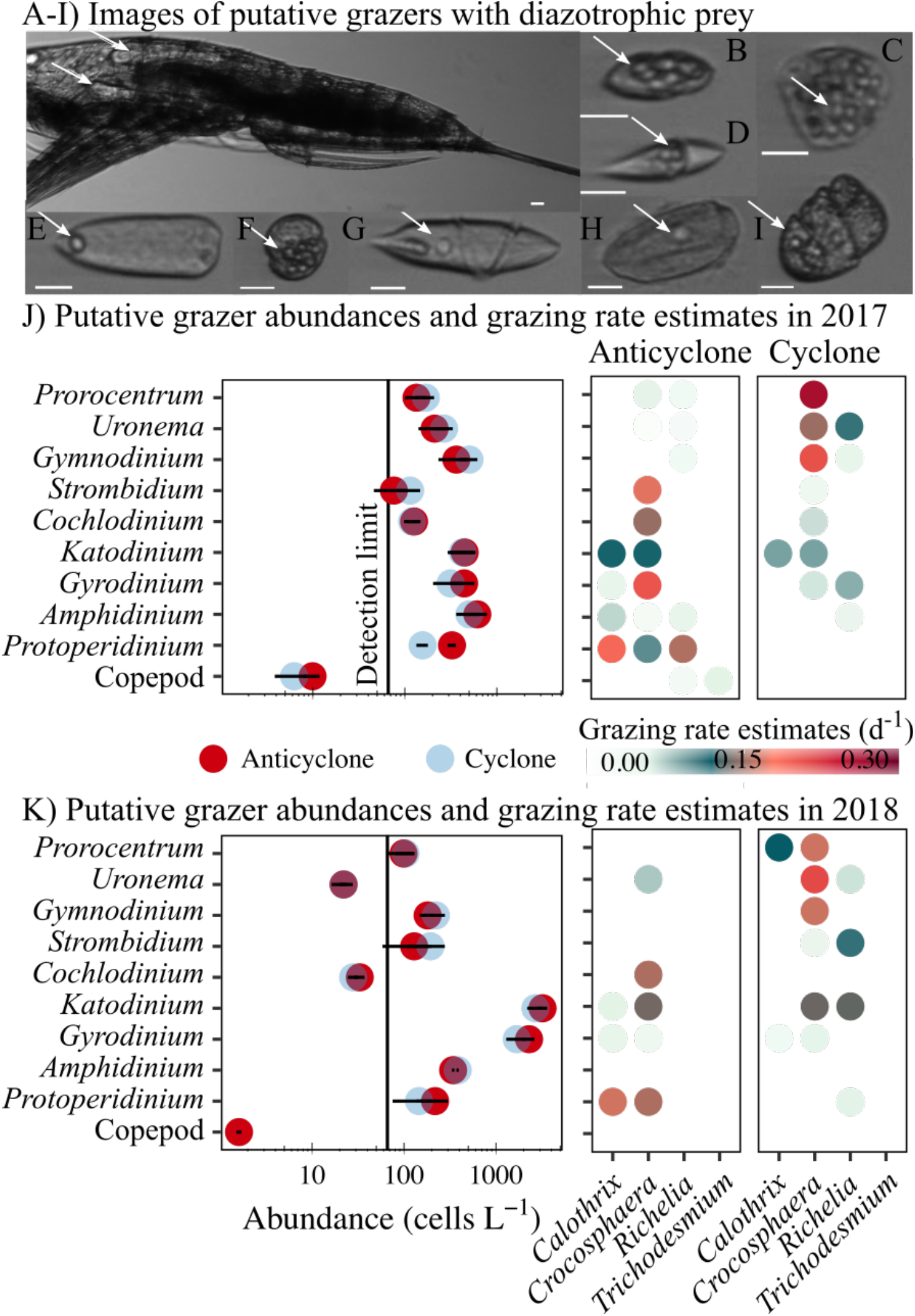
Evidence of diazotroph grazing in eddy cruises from images of co-occurring protists with ingested prey (A-I) and grazing rate estimates based on predator-prey dynamics (J-K). Grazing rates were predicted from protist abundances and interaction coefficients. Diazotrophic prey (indicated by arrows) were imaged inside copepods/nauplii (A), *Katodinium* (B), *Strombidium* (C), *Amphidinium* (D), *Prorocentrum* (E), *Gymnodinium* (F), *Gyrodinium* (H), *Uronema* (I), and *Cochlodinium* (J). Scale bars: 10 *µ*m.

The community structure of putative grazers varied across cruises and between eddies. Abundances of *Gymnodinium* (435 ± 100 cells L^-1^ in 2017 and 202 ± 29 cells L^-1^ in 2018), *Uronema* (239 ± 40 cells L^-1^ in 2017 and 22 ± 0 cells L^-1^ in 2018), *Cochlodinium* (123 ± 4 cells L^-1^ in 2017 and 31 ± 4 cells L^-1^ in 2018), *Amphidinium* (560 ± 78 cells L^-1^ in 2017 and 364 ± 29 cells L^-1^ in 2018), and copepods/nauplii (8 ± 3 cells L^-1^ in 2017 and 1.6 cells L^-1^ in 2018) were higher in the 2017 cruise, while abundances of *Strombidium* (162 ± 46 cells L^-1^ in 2018 and 95 ± 27 cells L^-1^ in 2017), *Katodinium* (2,958 ± 373 cells L^-1^ in 2018 and 438 ± 15 cells L^-1^ in 2017), and *Gyrodinium* (1,994 ± 430 cells L^-1^ in 2018 and 379 ± 90 cells L^-1^ in 2017) were higher in the 2018 cruise (Fig. 7J-K). Abundances of some taxa also varied with eddy polarity: *Prorocentrum*, *Uronema*, *Gymnodinium*, and *Strombidium* were generally more abundant in cyclones while *Gyrodinium*, *Amphidinium*, *Protoperidinium*, and copepods were more abundant in both anticyclones (Fig. 7). During the 2017 cruise, the community structure of grazers was also assessed by analyzing sequences (ribotags) from near the V4 region of the18S rRNA gene from metatranscriptomes (methods and results described in Supporting Information S5). These data likewise show different community structures of putative diazotroph grazers between eddies; for instance, the relative abundance of *Protoperidinium* ribotags was higher in the anticyclonic eddy, while *Strombidium* and *Cochlodinium* ribotags had higher relative abundances in the cyclone (Supporting Information Fig. S3).

Results of the predator-prey model provide estimates of grazing rates on specific diazotrophs during the two eddy cruises (Fig. 7). Maximum grazing rates ranged between 0.08-0.23 d^-1^ for heterotrophic dinoflagellates, between 0.12-0.22 d^-1^ for mixotrophic dinoflagellates, and 0.02-0.16 d^-1^ for ciliates (Fig. 7J-K). Predicted grazing rates varied between eddies according to grazer abundances, with the exception of *Cochlodinium*, *Katodinium*, and *Gyrodinium,* whose apparent abundances determined with the IFCb and metatranscriptome sequencing did not differ significantly (Fig. 7 J-K, Supporting Information S5). A few taxa had higher grazing rates in cyclones, including *Gymnodinium* (0.12 ± 0.04 d^-1^ in the cyclone and 0.001 ± 0.04 d^-1^ in the anticyclone) and *Prorocentrum* (0.18 ± 0.06 d^-1^ in the cyclone and 0.05 ± 0.06 d^-1^ in the anticyclone). IFCb analysis indicated that these appeared to graze mainly upon *Crocosphaera*, with a mean grazing pressure (the ratio of predicted grazing rate to growth rate) of 85 ± 46 % and higher grazing pressures in cyclonic eddies than anticyclonic eddies (Table 2). Grazers responsible for higher grazing rates in anticyclones included *Protoperidinium* (0.20 ± 0.04 d^-1^ in the anticyclone and 0.004 ± 0.006 d^-1^ in the cyclone), *Gyrodinium* (0.05 ± 0.06 d^-1^ in the anticyclone and 0.02 ± 0.03 d^-1^ in the cyclone), and *Amphidinium* (0.07 ± 0.004 d^-1^ in the anticyclone and 0.007 ± 0.01 d^-1^ in the cyclone). These grazers were associated with a high grazing pressure (mean 218 ± 13 %) in anticyclonic eddies upon *Calothrix* (Table 2). Predicted losses due to grazing of the large/endosymbiotic diazotroph *Richelia* were always below 0.1 d^−1^, resulting in a grazing pressure of 68 ± 45% in cyclones and 69% in anticyclones (Table 2).

## 5. Discussion

### 5.1 NFRs and diazotroph abundances at the mesoscale

Mesoscale eddies add variability to the biogeochemistry and microbial community structure in the NPSG (Barone et al., 2022; Barone et al., 2019), but the effects of eddies on N_2_ fixation are not fully understood. We sampled two pairs of eddies in spatial proximity, providing a unique opportunity to explore the effects of anticyclonic and cyclonic eddies on diazotrophy independent of seasonal changes. Since samples were collected in the vicinity of Station ALOHA, we also leveraged data from the HOT program (Karl & Church, 2014) to give historical and seasonal context to our observations.

NFRs in both anticyclone centers examined here were anomalously high relative to previous measurements at Station ALOHA. Surface (25 m) rates in these eddies exceeded all previous NFRs from the time-series (2012-2019), with values 3.7- and 1.8-fold above the highest previous HOT observations from the months of sampling (July and April, respectively) (Fig. 8). Between 2012 and 2019, 13 trackable eddies occupied Station ALOHA during a HOT cruise, yet no NFR measurements from these cruises occurred close to the eddy center (data not shown); hence, it is possible that we observed high NFRs in anticyclones because we specifically targeted the center of mesoscale eddies. In contrast, rates in both cyclones were within the range of previous observations. During the 2017 cruise, the range in depth-integrated NFRs between eddies, sampled 5 days and ∼2° latitude apart, exceeded the range in monthly mean NFRs from the time-series measurements (∼100-400 *µ*mol N m^−2^ d^−1^, Böttjer et al. 2017). Our results are similar to the studies of Fong et al. (2008) and Wilson et al. (2017), which both sampled anticyclonic eddies in the NPSG during July and observed anomalously high NFRs (surface rates of 8.6 and 10.9 nmol N L^−1^ d^−1^, respectively). The March-April 2018 cruise data also highlights that elevated NFRs in anticyclones are not necessarily restricted to warm (>25°C) summer months, as previously observed by Church et al. (2009).

**Figure 8:**
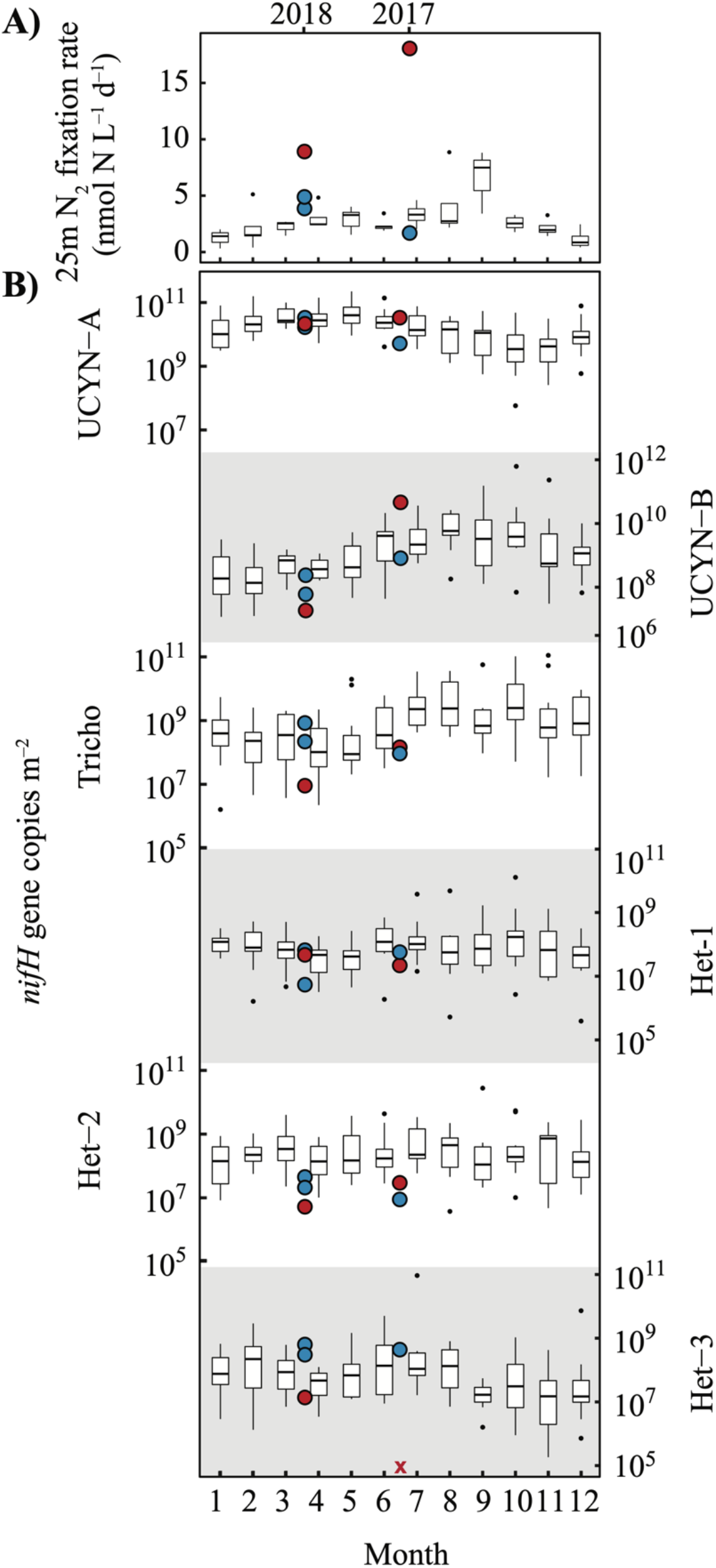
Eddy observations (cyclones [blue] and anticyclones [red]) of N_2_ fixation rates at 25 m depth (A, linear scale) and depth-integrated (0-125 m) *nifH* gene abundances (B, log scale) superimposed on historical HOT data. Boxplots illustrate ∼monthly measurements observed at Station ALOHA from 2012-2019 for N_2_ fixation rates (Böttjer et al., 2017) and from 2004-2017 for qPCR-derived *nifH* gene abundances (dataset doi: 10.5281/zenodo.4477269). Medians are shown as thick horizontal lines, 25-75% quantiles as boxes, the minimum and maximum values (up to 1.5 times the interquartile range) as whiskers, and outliers as black dots. Observations below detection limits are represented by colored ×. Shading is used to differentiate vertical panels.

The high NFRs observed in the 2017 and 2018 anticyclones appear to be driven by different diazotroph taxa. UCYN-A *nifH* gene abundances were the highest of all groups quantified during the spring (2018) cruise and UCYN-A and *Crocosphaera* (UCYN-B) were most abundant in the summer (2017) cruise (Fig. 3, Fig. 8). This is consistent with the known seasonal succession of *nifH* gene-based diazotroph abundances at Station ALOHA: abundances of *Trichodesmium*, *Crocosphaera*, and DDA *nifH* genes peak in the summer or early fall, while abundances of UCYN-A peak during the spring (Böttjer et al., 2014; Church et al., 2009). However, our only observation of both anomalously high diazotroph abundances and high anticyclone NFRs was during the 2017 cruise, when *Crocosphaera* abundances in the anticyclone exceeded all previous measurements in the region from the months of June and July (Fig. 8). The high abundances of *Crocosphaera* in the 2017 anticyclone were verified by independent cell counts via automated flow cytometry (Table 1). Although UCYN-A abundances were also high in the 2017 anticyclone, they only slightly exceeded the seasonal mean (Fig. 8). We infer that *Crocosphaera* likely played a major role in driving the high NFRs observed in the 2017 anticyclone. High *Crocosphaera* abundances have been recently linked to high NFRs in another anticyclonic eddy of the NPSG during July (Wilson et al., 2017).

It is less clear which diazotroph taxa drove the high NFRs observed in the 2018 anticyclone. In 2018, UCYN-A had the highest *nifH* gene abundances of all diazotrophs quantified; however, abundances were near the monthly mean and did not differ between eddies (Fig. 3, Fig. 8). No diazotroph taxa quantified via *nifH* gene abundances or automated flow cytometry had higher abundances in the anticyclonic eddy, where the highest rates were measured. There are several possible explanations for this apparent mismatch between diazotroph abundances and NFRs. NFRs vary as a function of both diazotroph abundance and cell-specific activity; hence, difference in rates between eddies could be due to differences in cell-specific activity rather than abundance. These differences were investigated for known cyanobacterial diazotrophs and are discussed in Section 5.2.1 below. Our ddPCR assays may also miss important diazotrophic taxa, such as non-cyanobacterial diazotrophs, whose importance in marine ecosystems are not well-understood (Moisander et al., 2017), or could misrepresent organismal abundances due to polyploidy (Sargent et al., 2016), low DNA extraction efficiency (Boström et al., 2004), or other methodological biases. Finally, the pattern observed in 2018 may be driven by patchiness of diazotroph communities (e.g. Robidart et al., 2014). We next examine evidence for eddy modulation of bottom-up, top-down, and physical control on diazotroph abundance and activity to better understand the mechanistic underpinnings of the observed patterns (Section 5.2). Finally, we address the challenges in generalizing the effects of eddies on N_2_ fixation due to confounding effects of season, diazotroph community structure, and eddy age/stage (Section 5.3).

### 5.2 Mechanisms leading to high NFRs in anticyclonic eddies

#### 5.2.1. Bottom-up control

Mesoscale eddies can modulate several factors that control diazotroph growth rates. Eddy dynamics can affect sea surface temperature via isopycnal uplift, resulting in cold core cyclones (McGillicuddy Jr & Robinson, 1997), and can affect light attenuation via nutrient injection and subsequent particle formation (Benitez-Nelson et al., 2007). Temperature and light both control cyanobacterial diazotroph growth rates (Supporting Information Table S3), and global NFRs correlate positively to both parameters (Luo et al., 2014). Eddies also influence nutrient availability (Hawco et al., 2021). Diazotrophs in the NPSG oscillate between P and Fe limitation (Letelier et al., 2019) and are able to coexist with non-diazotrophic phytoplankton under low N:P and N:Fe supply ratios (Ward et al., 2013). The uplift of isopycnals in cyclonic eddies can inject N, P, and Fe into the lower euphotic zone, where most of these nutrients are rapidly assimilated by the plankton community (McGillicuddy Jr & Robinson, 1997). Conversely, isopycnal downwelling in anticyclonic eddies can result in extreme N-depletion in surface waters, which has previously been used to explain observations of high NFRs in a NPSG anticyclone (Wilson et al., 2017).

In the present study, we have employed a suite of techniques that allowed us to address the potential limiting factors of diazotroph growth and modulation of these factors by mesoscale physical forcing. We found that strong differences in NFRs between the anticyclone and cyclone during the 2017 cruise may be partially driven by differences in PO_4_^3-^ availability. Surface PO_4_^3-^ concentrations and NFRs were significantly higher in the anticyclonic eddy than in the cyclonic eddy (Fig. 2). This pattern is consistent with a recent analysis of long-term trends at Station ALOHA documenting the depletion in PO_4_^3-^ concentrations in surface waters under low SLA (Barone et al., 2019). In our 2017 (summertime) cruise, surface PO_4_^3-^ in both 2017 eddies fell below 50 nmol L^−1^, the empirical threshold for P-limitation of diazotrophs in the NPSG reported by Letelier et al. (2019), and PO_4_^3-^ concentrations in the cyclone were below 30 nmol L^−1^, an apparent threshold for severe diazotroph P-limitation (Letelier et al., 2019). We hypothesize that the higher PO_4_^3-^ in the 2017 anticyclone partially relieved P-limitation, allowing for higher rates of diazotroph growth and N_2_ fixation by diazotrophic taxa, specifically *Crocosphaera*.

Eddy-driven reduction in P-limitation in the 2017 anticyclone is further supported by results from our ecophysiological model, patterns in nutrient stress marker gene expression, and previous studies. The ecophysiological model predicted that during the 2017 cruise, *Crocosphaera* was PO_4_^3-^-limited in the cyclone but not in the anticyclone, driving higher predicted growth rates in the anticyclone (Fig. 5). Indeed, in a complementary study that surveyed the 2017 eddies on the same expedition, *Crocosphaera* populations had higher intrinsic growth rates in the anticyclonic eddy than in the cyclonic eddy (Dugenne et al., 2020). Another study from the same 2017 expedition reported lower bulk alkaline phosphatase activity in the 2017 anticyclone (Harke et al., 2021), supporting the postulated reduced P-limitation there.

The metatranscriptome data provide additional evidence that PO_4_^3-^ availability helped drive the difference in *Crocosphaera* growth rates between eddies: all six *Crocosphaera* genes related to P acquisition and transport (commonly used as P-stress markers, Dyhrman & Haley, 2006; Pereira et al., 2016) had lower normalized transcript abundances in the 2017 anticyclone, consistent with a reduction in P-limitation relative to the cyclonic eddy (Fig. 6). We view the combined datasets presented here and in previous studies of the 2017 eddies as strong evidence that increased PO_4_^3-^ availability resulted in higher *Crocosphaera* growth rates in the anticyclone. This may ultimately help explain the higher *Crocosphaera nifH* gene abundances, biomass, and bulk NFRs observed in the anticyclone relative to the cyclone (Table 1, Fig. 3-4).

The higher concentrations of nutrients measured in 2018 did not seem to affect NFRs between cyclonic and anticyclonic eddies. Our ecophysiological model predicted that most diazotrophs were limited by low temperature, with the exception of DDAs, for which the thermal adaptation has not been tested in cultures. While data on the physiological adaptations of UCYN-A in culture are not available, a complementary study by Gradoville et al. (2021), which sampled eddies on the same 2018 expedition as the present study, reported no significant difference UCYN-A cell-specific NFRs between eddies, suggesting that bottom-up forcing did not differ between the 2018 eddies for this organism.

It is less likely that the differences in NFRs between cyclonic and anticyclonic eddies were driven by light, temperature, or dFe concentrations on either cruise. There were only small differences in near-surface light flux or temperatures between cyclones and anticyclones, except for the 2018 cruise, where lower daily-integrated light was observed in the anticyclone due to cloud cover (Fig. 2). Since higher NFRs were observed in the eddy with lower light levels, the opposite trend as expected (Luo et al., 2014), it is very unlikely that temperature or light drove the observed differences in NFRs between eddies, though they may help explain observed differences in NFRs between seasons (see Section 5.3) as well as variation in growth rates of distinct taxa (Fig. 5). Likewise, based on existing parameterization of Fe-uptake and Fe-stress marker genes expression, patterns in NFRs between eddies did not appear to be primarily driven by dFe. In the 2018 cruise, surface dFe concentrations were anomalously high in the cyclonic eddy, where lower NFRs were observed. In the 2017 cruise, dFe concentrations were slightly higher in the anticyclone, where rates were higher (Fig. 2). However, ecophysiological model results suggest that Fe was less limiting than PO_4_^3-^ at the surface for any diazotroph group on either cruise (Fig. 5). In addition, all surface dFe concentrations fell above the hypothesized Fe-limitation threshold for diazotrophs (0.13 nmol L^−1^), estimated from the empirical P-limitation threshold for diazotrophs in the NPSG (Letelier et al., 2019) coupled to diazotroph cell stoichiometry. There are caveats linked to testing the effect of iron concentration in culture and extrapolating these experiments to field populations, as described in Supporting Information S2. Nevertheless, expression patterns of Fe-stress marker genes (available for 2017 only) support our model results since most transcripts did not differ between cyclonic and anticyclonic eddies (Fig. 6). Collectively, these data are not consistent with an overriding control by Fe on NFR in our experiments.

We cannot exclude the possibility that there are other bottom-up control factors not analyzed here that could have potentially driven differences in diazotroph growth, and ultimately in bulk NFRs, between eddies. For example, some diazotrophs, such as *Trichodesmium,* use dissolved organic P (DOP) in P-depleted environments (Dyhrman et al., 2006; Orchard et al., 2010; White et al., 2010). Growth of symbiotic diazotrophs can also depend on the bottom-up control factors of their hosts, such as silica for the diatom hosts of *Richelia* and *Calothrix*. We did not report concentrations of DOP, silica, or any micronutrients other than dFe in this study due to a lack of data linking growth rates of cultured diazotrophs to the availability of these nutrients. As such, nutrients/micronutrients not assessed in our model could theoretically have driven differences in NFRs. In sum, of those potentially limiting abiotic factors measured, we find the strongest evidence of differential P limitation. While not exhaustive, our combined datasets, along with model results and long-term trends at station ALOHA (Barone et al. 2019), suggest that P may be the main abiotic factor driving the differences of NFRs in mesoscale eddies of the NPSG.

### 5.2.1. Top-down control

Little is known about the identity of diazotrophic grazers, or the extent to which grazing controls diazotroph abundances and NFRs in the NPSG (Landolfi et al., 2021). Early reports identified mesoplanktonic copepods grazing on *Trichodesmium* (Azimuddin et al., 2016; O’Neil et al., 1996; O’Neil, 1998), and later on UCYN-C (Hunt et al., 2016), UCYN-A (Scavotto et al., 2015), UCYN-B, Het-1, and Het-2 (Conroy et al., 2017). Recently, small nano/microplanktonic species (e.g. dinoflagellates and ciliates) were shown to prey on the large population of *Crocosphaera* in the NPSG (Dugenne et al., 2020). These small grazers generally dominate nano/microzooplankton biomass most of the year (Pasulka et al., 2013).

We identified numerous putative predators of diazotrophs using a predator-prey model and direct IFCb images of ingested prey: putative grazers spanned various sizes, from large copepods and nauplii imaged after having ingested *Trichodesmium* filaments, to nanoplanktonic ciliates and dinoflagellates mostly preying on *Crocosphaera* or *Calothrix*. Predicted rates varied between cyclonic and anticyclonic eddies (Fig. 7). Most of the grazers known to be mixotrophs (Jeong et al., 2010), such as *Prorocentrum*, *Gymnodinium*, and *Strombidium*, were more abundant in cyclones, while the heterotrophs *Protoperidinium*, *Amphidinium*, and *Katodinium* had higher abundances in anticyclones. Although understanding the drivers of grazer communities at the mesoscale is beyond the scope of this study, these differences in grazer abundances resulted in clear differences in predicted grazing rates upon specific diazotrophs in eddies of specific polarity and may help explain the observed patterns in diazotroph abundances and NFRs. During the 2017 cruise, Het-3 (*Calothrix*) *nifH* gene abundances measured within the mixed layer were 2-4 orders of magnitude greater in the cyclone then the anticyclone (Fig. 3) yet predicted growth rates did not differ between eddies (Fig. 5). Our predator-prey model predicts that rates of grazing upon *Calothrix* for the 2017 cruise were significantly higher in the anticyclone (0.18 ± 0.03 d^−1^, Table 2) than the cyclone (0.02 ± 0.004 d^−1^); thus, the observed differences in *Calothrix* abundance between eddies can simply be explained by differences in grazing pressures. In contrast, estimated rates of grazing upon *Crocosphaera* were higher in the cyclonic eddies for both cruises (Table 2). Therefore, the high abundance of *Crocosphaera* in the 2017 anticyclone appears driven by both enhanced growth rates and reduced grazing pressure, which ultimately helped to drive higher bulk NFRs (Fig. 3).

Results presented here and in previous studies (Dugenne et al., 2020; Landry et al., 2008; Wilson et al., 2017) show that grazing can be a significant loss term for small unicellular diazotrophs, but less so for larger diazotrophs such as *Richelia* and *Trichodesmium* (Table 2). Indeed, *Trichodesmium* has few known grazers (O’Neil & Roman, 1992), and losses are more often thought to be driven by viral lysis (Mulholland, 2007). While we did not directly assess viral lysis in this study, visual inspection of *Trichodesmium* filaments from the bloom at the edge of the 2017 cyclone (Fig. 4) suggests that *Trichodesmium* losses in the bloom may have been driven by programmed cell death or viral lysis (Berman-Frank et al., 2004; Hewson et al., 2004), as a majority of the filaments contained cells which were shrunken or lacking thylakoids and bloom filaments had significantly lower chlorophyll per cell than filaments imaged within the 2017 eddy centers (Supporting Information S4, Fig. S2). The fraction of dimly-fluorescing *Trichodesmium* filaments did not differ between eddies of specific polarity. We view this fraction as a proxy for senescence but other sources of diazotroph mortality, including viral lysis and programmed cell death, should be assessed directly in future studies.

### 5.2.2. Physical control

Buoyant plankton can be concentrated in areas of surface convergence, where cells are horizontally advected to the site of convergence and then acted upon by the opposing forces of downwelling and upward buoyancy (D’Asaro et al., 2018; Yoder et al., 1994). The buoyancy of diazotrophic taxa varies greatly: smaller diazotrophs (e.g. *Crocosphaera* and *Calothrix-Chaetoceros*) are negatively buoyant and sink (Bach et al., 2012; Tuo, 2015) while large diazotrophs (*Trichodesmium* and *Richelia* associations) can regulate their buoyancy, sometimes becoming positively buoyant and ascending the water column at high rates (Hoppe, 2013; Villareal & Carpenter, 2003; Villareal & Carpenter, 1989; Walsby, 1978). Elevated surface abundances of *Trichodesmium* and *Richelia* have been observed at numerous frontal features (Aldeco-Ramírez et al., 2009; Benavides et al., 2011; Guidi et al., 2012; Yoder et al., 1994), presumably due to frontal downwelling and strong positive cell buoyancy. In mesoscale eddies, downwelling can occur in intensifying anticyclones and weakening cyclones (McGillicuddy Jr, 2016), cyclones under constant wind (Gaube et al., 2015), and at the cyclonic side of eddy fronts (Mahadevan, 2016). The downwelling-induced physical accumulation of cells has been used to explain the occurrence of surface *Trichodesmium* slicks (Olson et al., 2015) and *Rhizosolenia* mats (Villareal & Carpenter, 1989) in eddies of both polarities.

The fine-scale distribution of diazotrophs across the pair of adjacent eddies measured in 2017 shows evidence for the physical accumulation of *Trichodesmium* and *Richelia* (Fig. 4). The highest *nifH* gene abundances of *Trichodesmium* and *Richelia* were observed near the surface at the cyclonic side of the front separating the two eddies, consistent with accumulation via surface downwelling caused by frontal dynamics. A surface bloom of *Trichodesmium* was also visually observed at the cyclonic side of the front, likely also driven by frontal downwelling. These results are consistent with a previous survey of *Trichodesmium* in an eddy dipole of the NPSG, which also reported maximal abundances at the front between eddies (Guidi et al., 2012).

Our observations are insufficient for determining the importance of physical accumulation on NFRs across the full eddy-eddy transect. NFR measurements were conducted in the center of each eddy but not on the cyclonic side of the eddy-eddy front where *Trichodesmium* and DDAs appeared to be physically accumulated during the 2017 cruise. Thus, while the eddy-induced physical accumulation of large diazotrophs likely explains elevated NFRs and abundances of large diazotrophs in some previous studies (e.g. Fong et al., 2008; Olson et al., 2015), our eddy observations were likely driven by differences in biological control—the balance of bottom-up and top-down control—between eddies.

### 5.3 Challenges in generalizing eddy effects on diazotrophs

Our finding of high NFRs in anticyclonic eddies agrees with other recent observations of high NFRs in anticyclones from many ocean regions (Fong et al., 2008; Holl et al., 2007; Liu et al., 2020; Löscher et al., 2016; Rahav et al., 2013; Wilson et al., 2017). However, there have also been reports of high diazotroph abundances associated with cyclonic eddies (Olson et al., 2015), and on a different cruise that sampled a cyclone-anticyclone eddy pair in the NPSG, surface NFRs were significantly higher in the cyclonic eddy than in the anticyclonic eddy (dataset doi: 10.5281/zenodo.5565560). At Station ALOHA, the correlation between NFR and SLA measured between 2012-2019 has remained non-significant (Spearman correlation, r=0.08, *p*-value=0.58, n=48, data not shown), yet high rates of N_2_ fixation are more prevalent in warm summer months often coincident with high SLA.

Despite this relative predictability, several taxa showed inconsistent patterns with eddy polarity during our cruises. Our data suggest that effects of eddies vary strongly among diazotroph taxa. Our ecophysiological model predicted bottom-up effects of mesoscale eddies to vary among diazotrophs, driven by the different adaptations of taxa to light, temperature, and nutrient concentrations. Likewise, eddy-specific grazing varied among diazotroph taxa, with some taxa being grazed upon more in cyclones and others being grazed upon more in anticyclones. Physical accumulation via frontal downwelling appeared to drive the elevated abundance of *Trichodesmium* and *Richelia* at the cyclonic side of the eddy-eddy front in 2017, but apparently did not affect the abundances of smaller, less buoyant taxa. Unfortunately, we were not able to assess eddy-driven mechanisms for some potentially important diazotroph taxa, including UCYN-A, whose growth and grazing could not be predicted due to the lack of culture data and the size constraints of IFCb imaging. Nevertheless, our results suggest that eddy effects on small diazotrophs like *Crocosphaera* are mostly due to biological effects (the net effects of growth and grazing), while eddy effects on large diazotrophs (*Trichodesmium* and DDAs) may be more affected by physical mechanisms.

Finally, changes in the properties of eddies through age and stage further complicates generalizing eddy effects. Mesoscale eddies are evolving features that intensify (with increasingly positive or negative SLA), reach a stable phase, and finally decay over the timescales of months (Sweeney et al., 2003). Though cyclonic and anticyclonic eddies are commonly associated with isopycnal uplift/depression and nutrient injection/dilution in the lower euphotic zone, respectively, in reality, vertical motions and nutrient fluxes vary with eddy age and stage (McGillicuddy Jr & Robinson, 1997). Our eddy observations, sampled over the timescales of days, represent snapshots of evolving features; we expect the eddy-induced mechanisms discussed above to vary through the lifetime of an eddy. At the time of sampling, both anticyclones were in a stable phase and both cyclones were in a weakening phase (Table S1), so these four observations are insufficient to deduce effects of eddy stage on NFRs.

## 6. Conclusions

Here we report anomalously high NFRs in two anticyclonic eddies in the NPSG. We coupled diverse datasets and ecological models to explore the mechanisms driving this pattern. Our analyses suggest that the 2017 anticyclone modulated bottom-up (via a reduction in PO_4_^3-^ limitation) and top-down (via eddy-specific grazing) control, allowing for the net accumulation of *Crocosphaera* cells, and ultimately, high NFRs. This mechanism may drive high NFRs in other anticyclonic eddies in the NPSG, where surface PO_4_^3-^ concentrations are significantly higher in anticyclones than in cyclones (Barone et al., 2019), especially during the summer months of peak *Crocosphaera* abundance. However, no drivers we tested appear to explain the results of our 2018 expedition, for which the high NFR observed in the anticyclonic eddy remains enigmatic. The biological and physical mechanisms through which eddies influence NFRs vary with season, diazotroph community composition, and eddy age/stage. Future efforts may benefit from assessing the ecological controls of uncultivated diazotrophs and from temporal measurements throughout the lifetime of eddies to better understand how eddies might affect successional patterns of diazotrophic communities and measured bulk NFRs.

## 7. Acknowledgements

This work was funded by the Simons Foundation (Award # 721252 to DMK, 721256 to AEW, 721223 to EFD, 721221 to MJC, 721244 to EVA, 721225 to STD, 329108 to SJ, and 724220 to JPZ) and expedition funding from the Schmidt Ocean Institute for R/V *Falkor* Cruise FK180310 in 2018. We are thankful to the chief scientists, including Tara Clemente (2017) and Steve Poulos (2018), as well as the captains and crew members of the eddy cruises. We are also grateful to Eric Shimabukuro, Ryan Tabata, and Tim Burrell for their help with field operations, as well as Katie Watkins-Brandt for valuable support with the IFCb on the 2017 expedition.

## 8. Data availability statement

NFRs and diazotroph abundances, measured by ddPCR and automated flow cytometry, along with environmental data and model output from the eddy cruises can be accessed at https://doi.org/10.5281/zenodo.6342202. Metatranscriptome sequences are available on the NCBI SRA under project numbers PRJNA596510 and PRJNA515070. NFRs and diazotroph *nifH* gene abundances from the Hawaii Ocean Time-series program can be accessed under the doi 10.5281/zenodo.3718435, 10.5281/zenodo.6341629, and 10.5281/zenodo.4477269.

## References

Aldeco-Ramírez, J., Monreal-Gómez, M., Signoret, M., Salas-de-León, D., & Hernández-Becerril, D. (2009). Occurrence of a subsurface anticyclonic eddy, fronts, and *Trichodesmium* spp. Ciencias Marinas, 35(4), 333–344.

Azimuddin, K. M., Hirai, J., Suzuki, S., Haider, M. N., Tachibana, A., Watanabe, K., Kitamura, M., Hashihama, F., Takahashi, K., & Hamasaki, K. (2016). Possible association of diazotrophs with marine zooplankton in the Pacific Ocean. MicrobiologyOpen, 5(6), 1016–1026.

Bach, L. T., Riebesell, U., Sett, S., Febiri, S., Rzepka, P., & Schulz, K. G. (2012). An approach for particle sinking velocity measurements in the 3–400 μm size range and considerations on the effect of temperature on sinking rates. Marine Biology, 159(8), 1853–1864.

Barone, B., Church, M. J., Dugenne, M., Hawco, N. J., Jahn, O., White, A. E., John, S. G., Follows, M. J., DeLong, E. F., & Karl, D. M. (2022). Biogeochemical dynamics in adjacent mesoscale eddies of opposite polarity. Global Biogeochemical Cycles, 36(2), e2021GB007115.

Barone, B., Coenen, A. R., Beckett, S. J., McGillicuddy, D. J., Weitz, J. S., & Karl, D. M. (2019). The ecological and biogeochemical state of the North Pacific Subtropical Gyre is linked to sea surface height. Journal of Marine Research, 77(2), 215–245.

Benavides, M., Agawin, N. S., Arístegui, J., Ferriol, P., & Stal, L. J. (2011). Nitrogen fixation by *Trichodesmium* and small diazotrophs in the subtropical northeast Atlantic. Aquatic Microbial Ecology, 65(1), 43–53.

Benitez-Nelson, C. R., Bidigare, R. R., Dickey, T. D., Landry, M. R., Leonard, C. L., Brown, S. L., Nencioli, F., Rii, Y. M., Maiti, K., & Becker, J. W. (2007). Mesoscale eddies drive increased silica export in the subtropical Pacific Ocean. Science, 316(5827), 1017–1021.

Berman-Frank, I., Bidle, K. D., Haramaty, L., & Falkowski, P. G. (2004). The demise of the marine cyanobacterium, *Trichodesmium* spp., via an autocatalyzed cell death pathway. Limnology and Oceanography, 49(4), 997–1005.

Boatman, T. G., Davey, P. A., Lawson, T., & Geider, R. J. (2018). The physiological cost of diazotrophy for *Trichodesmium erythraeum* IMS101. PloS One, 13(4), e0195638.

Boström, K. H., Simu, K., Hagström, Å., & Riemann, L. (2004). Optimization of DNA extraction for quantitative marine bacterioplankton community analysis. Limnology and Oceanography: Methods, 2(11), 365–373.

Böttjer, D., Dore, J. E., Karl, D. M., Letelier, R. M., Mahaffey, C., Wilson, S. T., Zehr, J., & Church, M. J. (2017). Temporal variability of nitrogen fixation and particulate nitrogen export at Station ALOHA. Limnology and Oceanography, 62, 200–216.

Böttjer, D., Karl, D. M., Letelier, R. M., Viviani, D. A., & Church, M. J. (2014). Experimental assessment of diazotroph responses to elevated seawater pCO_2_ in the North Pacific Subtropical Gyre. Global Biogeochem. Cycles, 28(6), 601–616.

Cheung, S., Nitanai, R., Tsurumoto, C., Endo, H., Nakaoka, S. i., Cheah, W., Lorda, J. F., Xia, X., Liu, H., & Suzuki, K. (2020). Physical forcing controls the basin-scale occurrence of nitrogen-fixing organisms in the North Pacific Ocean. Global Biogeochemical Cycles, e2019GB006452.

Church, M. J., Björkman, K. M., Karl, D. M., Saito, M. A., & Zehr, J. P. (2008). Regional distributions of nitrogen-fixing bacteria in the Pacific Ocean. Limnology and Oceanography, 53, 63–77.

Church, M. J., Jenkins, B. D., Karl, D. M., & Zehr, J. P. (2005). Vertical distributions of nitrogen-fixing phylotypes at Stn ALOHA in the oligotrophic North Pacific Ocean. Aquatic Microbial Ecology, 38(1), 3–14.

Church, M. J., Mahaffey, C., Letelier, R. M., Lukas, R., Zehr, J. P., & Karl, D. M. (2009). Physical forcing of nitrogen fixation and diazotroph community structure in the North Pacific subtropical gyre. Global Biogeochemical Cycles, 23(2), GB2020.

Church, M. J., Short, C. M., Jenkins, B. D., Karl, D. M., & Zehr, J. P. (2005). Temporal patterns of nitrogenase gene (*nifH*) expression in the oligotrophic North Pacific Ocean. Applied and Environmental Microbiology, 71(9), 5362–5370.

Conroy, B. J., Steinberg, D. K., Song, B., Kalmbach, A., Carpenter, E. J., & Foster, R. A. (2017). Mesozooplankton graze on cyanobacteria in the Amazon River plume and western tropical North Atlantic. Frontiers in Microbiology, 8, 1436.

D’Asaro, E. A., Shcherbina, A. Y., Klymak, J. M., Molemaker, J., Novelli, G., Guigand, C. M., Haza, A. C., Haus, B. K., Ryan, E. H., & Jacobs, G. A. (2018). Ocean convergence and the dispersion of flotsam. Proceedings of the National Academy of Sciences USA, 115(6), 1162–1167.

Davis, C. S., & McGillicuddy, D. J. (2006). Transatlantic abundance of the N_2_-fixing colonial cyanobacterium *Trichodesmium*. Science, 312(5779), 1517–1520.

de Boyer Montégut, C., Madec, G., Fischer, A. S., Lazar, A., & Iudicone, D. (2004). Mixed layer depth over the global ocean: An examination of profile data and a profile-based climatology. Journal of Geophysical Research: Oceans, 109, C12003.

Dugenne, M., Henderikx Freitas, F., Wilson, S. T., Karl, D. M., & White, A. E. (2020). Life and death of *Crocosphaera* sp. in the Pacific Ocean: Fine scale predator–prey dynamics. Limnology and Oceanography, 65, 2603–2617.

Dyhrman, S., Chappell, P., Haley, S., Moffett, J., Orchard, E., Waterbury, J., & Webb, E. (2006). Phosphonate utilization by the globally important marine diazotroph *Trichodesmium*. Nature, 439(7072), 68–71.

Dyhrman, S. T., & Haley, S. T. (2006). Phosphorus scavenging in the unicellular marine diazotroph *Crocosphaera watsonii*. Applied and Environmental Microbiology, 72(2), 1452–1458.

Farnelid, H., Andersson, A. F., Bertilsson, S., Al-Soud, W. A., Hansen, L. H., Sørensen, S., Steward, G. F., Hagström, Å., & Riemann, L. (2011). Nitrogenase gene amplicons from global marine surface waters are dominated by genes of non-cyanobacteria. PloS One, 6(4), e19223.

Ferrón, S., del Valle, D. A., Björkman, K. M., Quay, P. D., Church, M. J., & Karl, D. M. (2016). Application of membrane inlet mass spectrometry to measure aquatic gross primary production by the ^18^O *in vitro* method. Limnology and Oceanography: Methods, 14(9), 610–622.

Fitzsimmons, J. N., Hayes, C. T., Al-Subiai, S. N., Zhang, R., Morton, P. L., Weisend, R. E., Ascani, F., & Boyle, E. A. (2015). Daily to decadal variability of size-fractionated iron and iron-binding ligands at the Hawaii Ocean Time-series Station ALOHA. Geochimica et Cosmochimica Acta, 171, 303–324.

Follett, C. L., Dutkiewicz, S., Karl, D. M., Inomura, K., & Follows, M. J. (2018). Seasonal resource conditions favor a summertime increase in North Pacific diatom–diazotroph associations. The ISME journal, 12(6), 1543–1557.

Fong, A. A., Karl, D. M., Lukas, R., Letelier, R. M., Zehr, J. P., & Church, M. J. (2008). Nitrogen fixation in an anticyclonic eddy in the oligotrophic North Pacific Ocean. The ISME journal, 2(6), 663–676.

Foreman, R. K., Segura-Noguera, M., & Karl, D. M. (2016). Validation of Ti (III) as a reducing agent in the chemiluminescent determination of nitrate and nitrite in seawater. Marine Chemistry, 186, 83–89.

Foster, R., Subramaniam, A., Mahaffey, C., Carpenter, E., Capone, D., & Zehr, J. (2007). Influence of the Amazon River plume on distributions of free-living and symbiotic cyanobacteria in the western tropical north Atlantic Ocean. Limnology and Oceanography, 52(2), 517–532.

Gaube, P., Chelton, D. B., Samelson, R. M., Schlax, M. G., & O’Neill, L. W. (2015). Satellite observations of mesoscale eddy-induced Ekman pumping. Journal of Physical Oceanography, 45(1), 104–132.

Gifford, S. M., Becker, J. W., Sosa, O. A., Repeta, D. J., & DeLong, E. F. (2016). Quantitative transcriptomics reveals the growth-and nutrient-dependent response of a streamlined marine methylotroph to methanol and naturally occurring dissolved organic matter. MBio, 7(6), e01279–01216.

Gradoville, M. R., Cabello, A. M., Wilson, S. T., Turk-Kubo, K. A., Karl, D. M., & Zehr, J. P. (2021). Light and depth dependency of nitrogen fixation by the non-photosynthetic, symbiotic cyanobacterium UCYN-A. Environmental Microbiology, 23(8), 4518–4531.

Gradoville, M. R., Farnelid, H., White, A. E., Turk-Kubo, K. A., Stewart, B., Ribalet, F., Ferrón, S., Pinedo-Gonzalez, P., Armbrust, E. V., & Karl, D. M. (2020). Latitudinal constraints on the abundance and activity of the cyanobacterium UCYN-A and other marine diazotrophs in the North Pacific. Limnology and Oceanography, 65, 1858–1875. https://doi.org/10.1002/lno.11423

Guidi, L., Calil, P. H., Duhamel, S., Björkman, K. M., Doney, S. C., Jackson, G. A., Li, B., Church, M. J., Tozzi, S., & Kolber, Z. S. (2012). Does eddy-eddy interaction control surface phytoplankton distribution and carbon export in the North Pacific Subtropical Gyre? Journal of Geophysical Research: Biogeosciences, 117, G02024.

Harke, M. J., Frischkorn, K. R., Hennon, G. M., Haley, S. T., Barone, B., Karl, D. M., & Dyhrman, S. T. (2021). Microbial community transcriptional patterns vary in response to mesoscale forcing in the North Pacific Subtropical Gyre. Environmental Microbiology, 23(8), 4807–4822.

Hawco, N. J., Barone, B., Church, M. J., Babcock-Adams, L., Repeta, D. J., Wear, E. K., Foreman, R. K., Björkman, K. M., Bent, S., & Van Mooy, B. A. (2021). Iron depletion in the deep chlorophyll maximum: mesoscale eddies as natural iron fertilization experiments. Global Biogeochemical Cycles, 35(12), e2021GB007112.

Hewson, I., Govil, S. R., Capone, D. G., Carpenter, E. J., & Fuhrman, J. A. (2004). Evidence of *Trichodesmium* viral lysis and potential significance for biogeochemical cycling in the oligotrophic ocean. Aquatic Microbial Ecology, 36(1), 1–8.

Holl, C. M., Waite, A. M., Pesant, S., Thompson, P. A., & Montoya, J. P. (2007). Unicellular diazotrophy as a source of nitrogen to Leeuwin Current coastal eddies. Deep Sea Research Part II: Topical Studies in Oceanography, 54(8-10), 1045–1054.

Hoppe, K. S. (2013). The sinking rate and transparent exopolymer particle (TEP) production of Hemiaulus hauckii. Doctoral dissertation. University of Texas at Austin.

Hunt, B. P., Bonnet, S., Berthelot, H., Conroy, B. J., Foster, R. A., & Pagano, M. (2016). Contribution and pathways of diazotroph-derived nitrogen to zooplankton during the VAHINE mesocosm experiment in the oligotrophic New Caledonia lagoon. Biogeosciences, 13(10), 3131.

Jassby, A. D., & Platt, T. (1976). Mathematical formulation of the relationship between photosynthesis and light for phytoplankton. Limnology and Oceanography, 21(4), 540–547.

Jeong, H. J., Yoo, Y. D., Kim, J. S., Seong, K. A., Kang, N. S., & Kim, T. H. (2010). Growth, feeding and ecological roles of the mixotrophic and heterotrophic dinoflagellates in marine planktonic food webs. Ocean science journal, 45(2), 65–91.

Karl, D., Christian, J., Dore, J., Hebel, D., Letelier, R., Tupas, L., & Winn, C. (1996). Seasonal and interannual variability in primary production and particle flux at Station ALOHA. Deep Sea Research Part II: Topical Studies in Oceanography, 43(2-3), 539–568.

Karl, D., Letelier, R., Tupas, L., Dore, J., Christian, J., & Hebel, D. (1997). The role of nitrogen fixation in biogeochemical cycling in the subtropical North Pacific Ocean. Nature, 388, 533–538.

Karl, D., Michaels, A., Bergman, B., Capone, D., Carpenter, E., Letelier, R., Lipschultz, F., Paerl, H., Sigman, D., & Stal, L. (2002). Dinitrogen fixation in the world’s oceans. Biogeochemistry, 57(1), 47–98.

Karl, D. M., & Church, M. J. (2014). Microbial oceanography and the Hawaii Ocean Time-series programme. Nature Reviews: Microbiology, 12(10), 699–713.

Karl, D. M., Church, M. J., Dore, J. E., Letelier, R. M., & Mahaffey, C. (2012). Predictable and efficient carbon sequestration in the North Pacific Ocean supported by symbiotic nitrogen fixation. Proceedings of the National Academy of Sciences USA, 109(6), 1842–1849.

Karl, D. M., & Tien, G. (1992). MAGIC: A sensitive and precise method for measuring dissolved phosphorus in aquatic environments. Limnology and Oceanography, 37(1), 105–116.

Landolfi, A., Prowe, A., Pahlow, M., Somes, C. J., Chien, C.-T., Schartau, M., Koeve, W., & Oschlies, A. (2021). Can top-down controls expand the ecological niche of marine N_2_ fixers? Frontiers in Microbiology, 12, 690200.

Landry, M. R., Brown, S. L., Rii, Y. M., Selph, K. E., Bidigare, R. R., Yang, E. J., & Simmons, M. P. (2008). Depth-stratified phytoplankton dynamics in Cyclone Opal, a subtropical mesoscale eddy. Deep Sea Research Part II: Topical Studies in Oceanography, 55(10-13), 1348–1359.

Letelier, R. M., Björkman, K. M., Church, M. J., Hamilton, D. S., Mahowald, N. M., Scanza, R. A., Schneider, N., White, A. E., & Karl, D. M. (2019). Climate-driven oscillation of phosphorus and iron limitation in the North Pacific Subtropical Gyre. Proceedings of the National Academy of Sciences USA, 116(26), 12720–12728.

Letelier, R. M., White, A. E., Bidigare, R. R., Barone, B., Church, M. J., & Karl, D. M. (2017). Light absorption by phytoplankton in the North Pacific Subtropical Gyre. Limnology and Oceanography, 62(4), 1526–1540.

Liu, J., Zhou, L., Li, J., Lin, Y., Ke, Z., Zhao, C., Liu, H., Jiang, X., He, Y., & Tan, Y. (2020). Effect of mesoscale eddies on diazotroph community structure and nitrogen fixation rates in the South China Sea. Regional Studies in Marine Science, 35, 101106.

Löscher, C., Bourbonnais, A., Dekaezemacker, J., Charoenpong, C. N., Altabet, M. A., Bange, H. W., Czeschel, R., Hoffmann, C., & Schmitz, R. (2016). N_2_ fixation in eddies of the eastern tropical South Pacific Ocean. Biogeosciences, 13, 2889–2899.

Luo, E., Eppley, J. M., Romano, A. E., Mende, D. R., & DeLong, E. F. (2020). Double-stranded DNA virioplankton dynamics and reproductive strategies in the oligotrophic open ocean water column. The ISME journal, 14(5), 1304–1315.

Luo, Y.-W., Lima, I., Karl, D., Deutsch, C., & Doney, S. (2014). Data-based assessment of environmental controls on global marine nitrogen fixation. Biogeosciences, 11(3), 691–708.

Mahadevan, A. (2016). The impact of submesoscale physics on primary productivity of plankton. Annual Review of Marine Science, 8, 161–184.

McGillicuddy Jr, D. (2016). Mechanisms of physical-biological-biogeochemical interaction at the oceanic mesoscale. Annual Review of Marine Science, 8, 125–159.

McGillicuddy Jr, D., & Robinson, A. (1997). Eddy-induced nutrient supply and new production in the Sargasso Sea. Deep Sea Research Part I: Oceanographic Research Papers, 44(8), 1427–1450.

McGillicuddy Jr, D. J. (2014). Do *Trichodesmium* spp. populations in the North Atlantic export most of the nitrogen they fix? Global Biogeochemical Cycles, 28(2), 103–114.

Mohr, W., Großkopf, T., Wallace, D. W. R., & LaRoche, J. (2010). Methodological underestimation of oceanic nitrogen fixation rates. PloS One, 5(9), e12583.

Moisander, P. H., Beinart, R. A., Hewson, I., White, A. E., Johnson, K. S., Carlson, C. A., Montoya, J. P., & Zehr, J. P. (2010). Unicellular cyanobacterial distributions broaden the oceanic N_2_ fixation domain. Science, 327(5972), 1512–1514.

Moisander, P. H., Beinart, R. A., Voss, M., & Zehr, J. P. (2008). Diversity and abundance of diazotrophic microorganisms in the South China Sea during intermonsoon. The ISME journal, 2(9), 954–967.

Moisander, P. H., Benavides, M., Bonnet, S., Berman-Frank, I., White, A. E., & Riemann, L. (2017). Chasing after non-cyanobacterial nitrogen fixation in marine pelagic environments. Frontiers in Microbiology, 8, 1736.

Montoya, J. P., Voss, M., Kahler, P., & Capone, D. G. (1996). A simple, high-precision, high-sensitivity tracer assay for N_2_ fixation. Applied and Environmental Microbiology, 62(3), 986–993.

Mulholland, M. (2007). The fate of nitrogen fixed by diazotrophs in the ocean. Biogeosciences, 4(1), 37–51.

O’Neil, J., Metzler, P., & Glibert, P. (1996). Ingestion of ^15^N_2_-labelled *Trichodesmium* spp. and ammonium regeneration by the harpacticoid copepod *Macrosetella gracilis*. Marine Biology, 125(1), 89–96.

O’Neil, J. M. (1998). The colonial cyanobacterium *Trichodesmium* as a physical and nutritional substrate for the harpacticoid copepod *Macrosetella gracilis*. Journal of Plankton Research, 20(1), 43–59.

O’Neil, J. M., & Roman, M. R. (1992). Grazers and associated organisms of *Trichodesmium*. Marine pelagic cyanobacteria: Trichodesmium and other diazotrophs. Kluwer, 61–73.

Olson, E. M., McGillicuddy, D. J., Flierl, G. R., Davis, C. S., Dyhrman, S. T., & Waterbury, J. B. (2015). Mesoscale eddies and *Trichodesmium* spp. distributions in the southwestern North Atlantic. Journal of Geophysical Research: Oceans, 120(6), 4129–4150.

Orchard, E. D., Ammerman, J. W., Lomas, M. W., & Dyhrman, S. T. (2010). Dissolved inorganic and organic phosphorus uptake in *Trichodesmium* and the microbial community: The importance of phosphorus ester in the Sargasso Sea. Limnology and Oceanography, 55(3), 1390–1399.

Parks, D. H., Chuvochina, M., Waite, D. W., Rinke, C., Skarshewski, A., Chaumeil, P.-A., & Hugenholtz, P. (2018). A standardized bacterial taxonomy based on genome phylogeny substantially revises the tree of life. Nature Biotechnology, 36(10), 996–1004.

Pasulka, A. L., Landry, M. R., Taniguchi, D. A., Taylor, A. G., & Church, M. J. (2013). Temporal dynamics of phytoplankton and heterotrophic protists at station ALOHA. Deep Sea Research Part II: Topical Studies in Oceanography, 93, 44–57.

Pereira, N., Shilova, I. N., & Zehr, J. P. (2016). Molecular markers define progressing stages of phosphorus limitation in the nitrogen-fixing cyanobacterium, *Crocosphaera*. Journal of Phycology, 52(2), 274–282.

Pinedo-González, P., Hawco, N. J., Bundy, R. M., Armbrust, E. V., Follows, M. J., Cael, B., White, A. E., Ferrón, S., Karl, D. M., & John, S. G. (2020). Anthropogenic Asian aerosols provide Fe to the North Pacific Ocean. Proceedings of the National Academy of Sciences USA, 117(45), 27862–27868.

Poff, K. E., Leu, A. O., Eppley, J. M., Karl, D. M., & DeLong, E. F. (2021). Microbial dynamics of elevated carbon flux in the open ocean’s abyss. Proceedings of the National Academy of Sciences USA, 118(4), e2018269118.

Rahav, E., Herut, B., Stambler, N., Bar-Zeev, E., Mulholland, M. R., & Berman-Frank, I. (2013). Uncoupling between dinitrogen fixation and primary productivity in the eastern Mediterranean Sea. Journal of Geophysical Research: Biogeosciences, 118, 195–202.

Ribalet, F., Berthiaume, C., Hynes, A., Swalwell, J., Carlson, M., Clayton, S., Hennon, G., Poirier, C., Shimabukuro, E., White, A., & Armbrust, E. V. (2019). SeaFlow data v1, high-resolution abundance, size and biomass of small phytoplankton in the North Pacific. Scientific data, 6(1), 1–8.

Rii, Y. M., Peoples, L. M., Karl, D. M., & Church, M. J. (2021). Seasonality and episodic variation in picoeukaryote diversity and structure reveal community resilience to disturbances in the North Pacific Subtropical Gyre. Limnology and Oceanography.

Robidart, J. C., Church, M. J., Ryan, J. P., Ascani, F., Wilson, S. T., Bombar, D., Marin III, R., Richards, K. J., Karl, D. M., Scholin, C. A., & Zehr, J. P. (2014). Ecogenomic sensor reveals controls on N_2_-fixing microorganisms in the North Pacific Ocean. The ISME journal, 8(6), 1175.

Sargent, E. C., Hitchcock, A., Johansson, S. A., Langlois, R., Moore, C. M., LaRoche, J., Poulton, A. J., & Bibby, T. S. (2016). Evidence for polyploidy in the globally important diazotroph *Trichodesmium*. FEMS Microbiology Letters, 363(21), fnw244.

Scavotto, R. E., Dziallas, C., Bentzon-Tilia, M., Riemann, L., & Moisander, P. H. (2015). Nitrogen-fixing bacteria associated with copepods in coastal waters of the North Atlantic Ocean. Environmental Microbiology, 17(10), 3754–3765.

Snow, J. T., Polyviou, D., Skipp, P., Chrismas, N. A., Hitchcock, A., Geider, R., Moore, C. M., & Bibby, T. S. (2015). Quantifying integrated proteomic responses to iron stress in the globally important marine diazotroph *Trichodesmium*. PloS One, 10(11), e0142626.

Sosik, H. M., & Olson, R. J. (2007). Automated taxonomic classification of phytoplankton sampled with imaging-in-flow cytometry. Limnology and Oceanography: Methods, 5(6), 204–216.

Stenegren, M. (2020). Significance of N2 fixing planktonic symbioses for open ocean ecosystems [Doctoral dissertation, Stockholm University].

Stukel, M. R., Coles, V. J., Brooks, M., & Hood, R. R. (2014). Top-down, bottom-up and physical controls on diatom-diazotroph assemblage growth in the Amazon River plume. Biogeosciences, 11(12), 3259–3278.

Sweeney, E. N., McGillicuddy Jr, D. J., & Buesseler, K. O. (2003). Biogeochemical impacts due to mesoscale eddy activity in the Sargasso Sea as measured at the Bermuda Atlantic Time-series Study (BATS). Deep Sea Research Part II: Topical Studies in Oceanography, 50(22-26), 3017–3039.

Taboada, F. G., Gil, R. G., Höfer, J., González, S., & Anadón, R. (2010). *Trichodesmium* spp. population structure in the eastern North Atlantic subtropical gyre. Deep Sea Research Part I: Oceanographic Research Papers, 57(1), 65–77.

Tibshirani, R., Bien, J., Friedman, J., Hastie, T., Simon, N., Taylor, J., & Tibshirani, R. J. (2012). Strong rules for discarding predictors in lasso-type problems. *Journal of the Royal Statistical Society*, Series B (Statistical Methodology), 74(2), 245–266.

Tuo, S. (2015). The distributions and the mechanisms of the cyanobionts and their host diatoms in the northern South China Sea and the Kuroshio [Doctoral dissertation, National Sun Yat-sen University].

Villareal, T., & Carpenter, E. (2003). Buoyancy regulation and the potential for vertical migration in the oceanic cyanobacterium *Trichodesmium*. Microbial Ecology, 45(1), 1–10.

Villareal, T. A., & Carpenter, E. J. (1989). Nitrogen fixation, suspension characteristics, and chemical composition of *Rhizosolenia* mats in the central North Pacific gyre. Biological Oceanography, 6(3-4), 327–345.

Walsby, A. (1978). The properties and buoyancy-providing role of gas vacuoles in *Trichodesmium* Ehrenberg. British Phycological Journal, 13(2), 103–116.

Ward, B. A., Dutkiewicz, S., Moore, C. M., & Follows, M. J. (2013). Iron, phosphorus, and nitrogen supply ratios define the biogeography of nitrogen fixation. Limnology and Oceanography, 58(6), 2059–2075.

Webb, E. A., Ehrenreich, I. M., Brown, S. L., Valois, F. W., & Waterbury, J. B. (2009). Phenotypic and genotypic characterization of multiple strains of the diazotrophic cyanobacterium, *Crocosphaera watsonii*, isolated from the open ocean. Environmental Microbiology, 11(2), 338–348.

White, A. E., Granger, J., Selden, C., Gradoville, M. R., Potts, L., Bourbonnais, A., Fulweiler, R. W., Knapp, A. N., Mohr, W., & Moisander, P. H. (2020). A critical review of the ^15^N_2_ tracer method to measure diazotrophic production in pelagic ecosystems. Limnology and Oceanography: Methods, 18(4), 129–147.

White, A. E., Karl, D. M., Bjorkman, K. M., Beversdorf, L. J., & Letelier, R. M. (2010). Production of organic matter by *Trichodesmium* IMS101 as a function of phosphorus source. Limnology and Oceanography, 55(4), 1755–1767. <Go to ISI>://WOS:000283657000023

Wilson, S. T., Aylward, F. O., Ribalet, F., Barone, B., Casey, J. R., Connell, P. E., Eppley, J. M., Ferrón, S., Fitzsimmons, J. N., & Hayes, C. T. (2017). Coordinated regulation of growth, activity and transcription in natural populations of the unicellular nitrogen-fixing cyanobacterium *Crocosphaera*. Nature Microbiology, 2(9), 17118.

Wilson, S. T., Böttjer, D., Church, M. J., & Karl, D. M. (2012). Comparative assessment of nitrogen fixation methodologies, conducted in the oligotrophic North Pacific Ocean. Applied and Environmental Microbiology, 78(18), 6516–6523.

Wozniak, B., Dera, J., Ficek, D., Majchrowski, R., Ostrowska, M., & Kaczmarek, S. (2003). Modelling light and photosynthesis in the marine environment. Oceanologia, 45(2), 171–245.

Yoder, J. A., Ackleson, S. G., Barber, R. T., Flament, P., & Balch, W. M. (1994). A line in the sea. Nature, 371(6499), 689–692.

Zeev, E. B., Yogev, T., Man-Aharonovich, D., Kress, N., Herut, B., Béja, O., & Berman-Frank, I. (2008). Seasonal dynamics of the endosymbiotic, nitrogen-fixing cyanobacterium *Richelia intracellularis* in the eastern Mediterranean Sea. The ISME journal, 2(9), 911–923.

Zehr, J. P., Waterbury, J. B., Turner, P. J., Montoya, J. P., Omoregie, E., Steward, G. F., Hansen, A., & Karl, D. M. (2001). Unicellular cyanobacteria fix N_2_ in the subtropical North Pacific Ocean. Nature, 412(6847), 635–637.

