## Supporting Information for "Nitrogen fixation in mesoscale eddies of the North Pacific Subtropical Gyre: patterns and mechanisms"

*Global Biogeochemical Cycles*

### Contents of this file

Text S1 to S5
Figures S1 to S3
Tables S1 and S3

### Additional Supporting Information (Files uploaded separately)

Captions for Tables S2 and S4

**Text S1: Characteristics of the eddies sampled during the eddy cruises**

The pairs of cyclonic and anticyclonic eddies sampled in 2017 and 2018 were identified via satellite-derived sea level anomalies (SLA) distributed by the Copernicus Marine Service (marine.copernicus.eu). Daily SLA maps were corrected for the interannual trend and seasonal cycle, and the targeted eddies were tracked back in time following the procedure described by (Barone et al., 2022). Briefly, eddy centers were tracked by identifying maxima and minima of corrected SLA in a window of 1.5° (in latitude and longitude) centered around the location of the eddy center on the previous day. The age, amplitude, and stage of each eddy at the time of sampling are based on the time-series of daily corrected SLA; the latter was determined by the slope of the time-series (ΔSLA/Δt) within 10 d before/after the sampling periods. The position of the front between pairs of eddies was determined using daily maps of finite size Lyapunov Exponent (FSLE), a measure of the divergence of parcels of waters in close proximity, distributed by AVISO (www.aviso.altimetry.fr). We followed the classification of Guo et al. (2019) for the North Pacific Ocean to locate frontal stations, defined as having elevated FSLE (>0.064 $d^{-1}$) and low absolute SLA (<0.048 m). The mixed layer depth (MLD) was determined in each eddy as the shallowest depth at which the potential density was at least 0.03 kg $m^{-3}$ greater than the potential density at 10 m (de Boyer Montégut et al., 2004).

**Table S1:** Characteristics of the pairs of cyclonic and anticyclonic eddies sampled during the eddy cruises

| | | Anticyclones | | | | | Cyclones | | | | |
|---|---|---|---|---|---|---|---|---|---|---|---|
| Year | Season | Phase | Age (d) | SLA (m) | ΔSLA/Δt (cm $d^{-1}$) | MLD (m) | Phase | Age (d) | SLA(m) | ΔSLA/Δt (cm $d^{-1}$) | MLD (m) |
| 2017 | Summer | stable | 83 | 0.26 | $0.04^{\dagger}$ | 23.9 ± 5.6 | weakening | 246 | -0.14 | 0.11 | 30.6 ± 8.6 |
| 2018 | Spring | stable | 95 | 0.21 | $0.02^{\dagger}$ | 50.3 ± 17.8 | weakening | 212 | -0.14 | 0.26 | 17.8 ± 5.6 |

$^{\dagger}$: Not significant

**Table S2:** Raw data and detection limit calculations for $N_2$ fixation rate (NFR) measurements on the two eddy cruises. The contribution of each source of error to the total uncertainty is provided (Montoya et al., 1996). The contribution of error associated with atom% $N_2$ is not included since only one MIMS sample was analyzed per batch of $^{15}N_2$-enriched seawater on all cruises. δNFR/ δX represent the partial derivative of the NFR with respect to each parameter, evaluated using respective means and standard deviations. Error Contribution and % Total Error represent the absolute and relative error associated with each parameter. The total uncertainty associated with each measurement is termed the "Minimal Quantifiable Rate" and calculated using standard propagation of error. Limits of detection (LOD) were calculated by setting the change in atom % $^{15}N$ of particulate N equal to 0.00146 atom %. NFR units are nmol N $L^{-1}$ $d^{-1}$. $T_{zero}$ natural abundance samples ($A_{PN0}$) were not collected during the 2017 cruise; hence, mean and standard deviation from the corresponding depth in 2018 were used for calculations.

*See "Supporting Information Tables S2 and S4" excel spreadsheet*

**Text S2: Ecophysiological model uncertainties, assumptions, and caveats**

We explored how mesoscale eddies affected the growth of cyanobacterial diazotrophs using a standard ecophysiological model (Follett et al., 2018; McGillicuddy Jr, 2014; Stukel et al., 2014). The model is based on parameters compiled from culture studies testing the effect of light, temperature, and nutrient concentrations on the intrinsic growth rates of *Crocosphaera*, *Trichodesmium*, and DDA symbionts. The compilation of diazotroph ecophysiological parameters is presented in Table S3, including the references of each study used. Since parameters may vary depending on the model formulation (see Tian and Chen (2006) for a review of the common formulations used to model phytoplankton growth), we consistently fitted the same formulation to each diazotrophic population using the nonlinear least-squares R function 'nls' and reported the predicted parameter with its standard deviation. In our model, we favored the formulation that includes the least parameters, unless additional parameters had been reported for multiple taxa, such as the light compensation coefficient in light response curves (Boatman et al., 2017; N. S. Garcia et al., 2013).

We used the environmental gradients of temperature (T), daily average instantaneous PAR (E), and nutrient concentrations to predict diazotroph growth rates and their uncertainties, $\mu(T,E, PO_4^{3-},dFe) \pm sd$, across eddy centers (Table 2). Standard deviations were used to propagate uncertainties to predictions of diazotroph growth rates. Our model assumes that phosphate ($PO_4^{3-}$) and dissolved iron (dFe) are the main nutrients that limit diazotroph growth rates (Sohm et al., 2011), and that their intrinsic growth rates are eventually limited by the resource in shortest supply. This assumption follows the classical Liebig's law of the minimum, with resources extended to light, which is required by cyanobacterial diazotrophs, and temperature, as a narrow temperature range (20-30°C) is thought to limit the global distribution of *Crocosphaera*, *Trichodesmium*, and DDAs (Tang et al., 2020). Note that in our study, temperatures never reached the thermal optimum for any diazotroph taxon examined (Fig. 2), but alternative models should be used in order to express the deleterious effect of temperatures above the thermal optimum if necessary. Since the thermal niche of DDAs has not been characterized in cultures, the effect of temperature on

*Richelia* and *Calothrix* growth rates was excluded. Similar ecological models have predicted the growth of DDAs in the Atlantic (Stukel et al., 2014) and the NPSG (Follett et al., 2018) without accounting for temperature.

There are multiple caveats to our modeling approach (numbered 1-5 below). We lack some ecophysiological parameters for cyanobacterial diazotrophs (1) and there is potential interdependency among parameters (2). Other caveats are linked to the differences between lab isolates and field populations (3), as well as between the in situ and tested conditions (4). Both differences should be minimized in order to scale up lab studies to the field. Finally, the model does not account for the role of additional nutrients, certain transient or symbiotic partnerships, and foraging behavior involved in diazotroph resource acquisition (5). Each caveat is detailed below:

**1) Lack of ecophysiological parameters:** Cultures of cyanobacterial diazotrophs have been used to inform how the growth limitations of individual taxa may differ amongst strains/ecotypes (i.e. physiological acclimation or plasticity) and/or over time (i.e. evolutionary adaptations). Both short- and long-term differences result from the genetic diversity of field populations and can be characterized in controlled conditions if researchers can isolate, maintain (over a few to hundreds of generations), and measure the response of specific strains to changes in abiotic factors. Culture studies focusing on marine cyanobacterial diazotrophs have been heavily biased towards *Trichodesmium erythraeum* and *Crocosphaera watsonii.* Symbiotic organisms are notably difficult to maintain in culture, hence information on DDAs, as well as UCYN-A and its eukaryotic host, have been limited (Caputo et al., 2018; Foster et al., 2010; Pyle et al., 2020; Villareal, 1989). To our knowledge, culture-based measurements of UCYN-A growth rates in response to light, temperature, or nutrient concentrations have not been reported; thus, we were not able to include UCYN-A in the ecophysiological model. Similarly, the effects of temperature and nutrient concentrations on the diazotrophic symbionts of DDAs (*Richelia* and *Calothrix*) have not been reported. We used the unique $PO_4^{3-}$ and dFe half-saturation coefficients (without uncertainties) described in Follett et al. (2018) for both DDA symbionts.

**2) Interdependency among parameters:** The ecophysiological traits of cyanobacterial diazotrophs, as in many phytoplankton taxa, are interdependent (Litchman et al., 2012). For instance, the light saturation parameter ($E_k$) is typically inversely correlated to $\mu_{max}$, the theoretical maximum growth rate (Edwards et al., 2015). The latter affects the nutrient half-saturation constants (i.e. the nutrient concentration at which the measured growth rate equals half the maximum growth rate) and increases exponentially with temperature (Eppley, 1972). To account for the trade-offs between $\mu_{max}$ and the other parameters, we normalized the growth rate estimates measured in cultures to their relative maximum. Doing so limited the effect of interactions and feedback loops among bottom-up controls (Boatman et al., 2018; Qu et al., 2019) and allowed us to compare studies performed under various experimental conditions that may prevent the culture from growing optimally. For example, Fu et al. (2005) compared the half-saturation constant for $PO_4^{3-}$ uptake of two strains of *Trichodesmium erythraeum* (IMS101 and GBRTRLI101) maintained at 25°C and an irradiance of 45 ±2 μmol quanta $m^{-2}$ $s^{-1}$. In their study, the strain GBRTRLI101 grew significantly slower than IMS101 under P-replete conditions, reaching a maximum growth rate of 0.18±0.01 $d^{-1}$, compared to 0.25±0.01 $d^{-1}$. Due to the differences in $\mu_{max}$, the authors reported a half-saturation constant ($k_{PO_4^{3-}}$) twice as large for *Trichodesmium* GBRTRLI101. Other studies, focusing on the effect of temperature on this strain, have shown that under P-replete conditions, *Trichodesmium* GBRTRLI101 may grow at rates exceeding 0.3 $d^{-1}$ (Fu et al., 2005; Qu et al., 2019). When normalized to their respective maximum, the growth kinetics of the strains IMS101 and GBRTRLI101 closely aligned with one another, and $k_{PO_4^{3-}}$ became non-significantly different (data not shown). In the present study, $\mu_{max}$ correspond to the maximum value reported across all literature values.

Perhaps more puzzling is the link between P- and Fe-uptake, both constrained by specific half-saturation constants. One mechanism linking Fe and P physiology is the P-stress induced production of enzymes with Fe co-factors, such as alkaline phosphatase (Rouco et al., 2018). Garcia et al. (2015) provided critical insight into the dependency of nutritional status in marine diazotrophs. They showed that $PO_4^{3-}$ uptake was significantly affected by the ambient concentration of bioavailable iron, which led to optimal rates of growth and $N_2$ fixation of *Crocosphaera* and *Trichodesmium* under co-stress conditions. There are a number of hypotheses that address this paradox, but at least in *Trichodesmium* (which reduces $CO_2$ and $N_2$ simultaneously), there seems to be a clear role of alternative electron transport pathways that require less iron than PSI while fueling the nitrogenase, as well as cell size reduction, as highlighted in recent metatransciptomic and metaproteomic studies (Walworth et al., 2016). The low $PO_4^{3-}$ and dFe concentrations used to simulate co-stress conditions in culture are representative of the NPSG, suggesting that the half-saturation for *Crocosphaera* and *Trichodesmium* phosphate uptake reported in Table S3 is adapted to our study site. Still, we cannot exclude the possibility that our model results are oversimplified by not accounting for nutrient colimitation effects in other organisms or for other interdependencies among model parameters.

**3) Differences between laboratory isolates and field populations:** Controlled studies of laboratory isolates have shown that diazotroph strains present unique ecophysiological plasticity. This plasticity seems to be primarily linked to phenotypic traits, rather than the geographic origin of each isolate (Boyd et al., 2013; Fu et al., 2014; Hynes, 2009; Webb et al., 2009). The potential for strain-specific responses of diazotroph taxa to light, temperature, and nutrient concentrations is a limitation to our approach.

Most laboratory isolates of *Crocosphaera* can be separated into two phenotypes (i.e. small- and large-size) with a distinction in thermal tolerance (Webb et al. (2009): large-size isolates being more tolerant to +/- temperature anomalies), maximum growth rates (N. Garcia et al. (2013): large-size isolates achieving higher maximum growth rates), and presumably, in their ability to scavenge phosphorus and iron, as suggested from the sequencing analysis of their genome (Bench et al. (2013): large-size isolates having a larger genome including additional genes linked to scavenging). The effects of $PO_4^{3-}$ and dFe concentrations on *Crocosphaera* growth rates have only been tested on large-sized *Crocosphaera watsonii* WH0003 (Garcia et al., 2015) and intermediate-sized *Crocosphaera watsonii* WH8501 (Jacq et al., 2014), respectively. In the present study, since *Crocosphaera* appeared primarily P-limited within the mixed layer, as indicated by the metatranscriptomic analyses of stress marker genes reflecting all phenotypes, we only predicted growth rates for the large-sized cells, where the half-saturation for P-uptake has been characterized in culture (Table S3). In comparison, the light saturation parameter of light-growth responses reported by N. Garcia et al. (2013) were relatively conserved across *Crocosphaera* isolates and were comparable to in situ estimates derived from a field population (Dugenne et al., 2020). We note that both small- and large-sized *Crocosphaera* populations were observed in our study, so modeling the response of large-sized *Crocosphaera* only may have biased our results.

The distribution of *Trichodesmium* field populations is linked to niche-adapted species and colony morphology, but only a subset of *Trichodesmium* species have been studied in culture (Hynes et al., 2012). Of *Trichodesmium* colonies collected at Station ALOHA, >99% of sequences belonged to Clade I, which includes the isolate *T. thiebautii,* and less than 1% of sequences belonged to Clade III, which includes *T. erythraeum* (the species most often used in culture-based studies) (Gradoville et al., 2017).(Rouco, Haley, Alexander, et al., 2016). The recent discovery of new non-diazotrophic *Trichodesmium* species, for which there are no isolates, further exacerbates the challenges associated with comparing cultures to field populations (Delmont, 2021). In the present study, we used growth parameters from studies on *T. erythraeum* strain IMS101, since responses of *T. thiebautii* to light, temperature, and nutrient concentrations have not been characterized in culture. However, we note that the ecophysiological adaptations of a field population likely dominated by *Trichodesmium* Clade I may diverge from that of *T. erythraeum* strain IMS101, since species from Clade I and Clade III appear to respond differently to environmental drivers such as $pCO_2$ (Hutchins et al., 2013) and contain different phosphorus-regulated genes (Orchard et al., 2003). We also note that photo-inhibition have been previously observed for *Trichodesmium* IMS101 at light intensity >1200 µmol quanta $m^{-2}$ $s^{-1}$ (Breitbarth et al., 2008), but the average light intensity never exceeded 1200 µmol quanta $m^{-2}$ $s^{-1}$ in our study, hence we did not account for photo-inhibition when parameterizing the effect of light intensity on diazotroph growth rates.

**4) Differences between in situ and culture conditions:** Ideally, the conditions tested in cultures should be similar to in situ abiotic factors in order to minimize the caveats of ecophysiological models. In cultures, the effect of phosphorus and iron limitation have been tested using simple molecules (e.g. $PO_4^{3-}$ or EDTA), which do not reflect the variety of dissolved compounds present in the environment. Furthermore, the concentrations tested are not always representative of in situ concentrations, particularly for iron. In the environment, dissolved iron is nearly entirely (80-90%) associated with organic ligands (a fraction of which being high Fe-affinity siderophores synthesized by bacteria to acquire Fe) of largely unknown identity, that cannot be reproduced in the lab. Free inorganic iron (Fe') is the only form measured in bioassays and is assumed to be the only bioavailable source of iron for cultures. At Station ALOHA, the depth profile of dFe concentration ranges between 0.06 and 0.15 nmol $L^{-1}$ and derived Fe' are predicted to be ~three orders of magnitude lower (0.05-0.14 pmol $L^{-1}$) (Bundy et al., 2018). In contrast, the Fe' tested in diazotroph cultures generally range between 20-6000 pmol $L^{-1}$ (Boatman et al., 2018). The kinetic curves are therefore poorly constrained under ultraoligotrophic Fe conditions and it is unclear whether a minimum quota should be considered and parameterized to predict diazotroph growth rates accurately. The (sometimes axenic) laboratory isolates also represent simplified biosystems, in which the influence of several life-history traits affecting diazotroph resource acquisition/storage have not been clearly tested. Hence, of all the parameters compiled in our study, the half-saturation constant for phosphorus and iron uptake may be the most biased. Perhaps all these biases can explain the conflicting studies comparing the affinity of *Trichodesmium* and *Crocosphaera* for dissolved iron (Jacq et al., 2014; Saito et al., 2011).

**5) Additional nutrients, partnerships, and behavior:** Under simplified conditions, the complex life-history traits of laboratory isolates are often overlooked. These include the aggregate/colonial stages of *Crocosphaera* and *Trichodesmium*, the latter involved in active vertical migration to orchestrate nutrient acquisition at depth and fixation of $CO_2$ and $N_2$-fixation closer to the surface (Foster et al., 2013; Hewson et al., 2009; Villareal & Carpenter, 2003; White et al., 2006), or the free-living stage of *Richelia* and *Calothrix* (Villareal, 1990; Villareal & Carpenter, 1989) which have been reported under high nitrate concentrations (Tuo et al., 2017; Villareal, 1990; Villareal & Carpenter, 1989). In addition, a number of partnerships have been reported in the field, including the *Trichodesmium* microbiome (Hmelo et al., 2012;

Rouco, Haley, & Dyhrman, 2016), *Trichodesmium* associated with *Calothrix* (Momper et al., 2015), *Crocosphaera* associated with *Climacodium* (Foster et al., 2011), and DDAs associated with pennate diatoms (Villareal et al., 2012). Some of these associations have been specifically implicated in resource acquisition (Basu et al., 2019; Frischkorn et al., 2018; Frischkorn et al., 2017; Gradoville et al., 2017). Thus, our model could have misrepresented the responses of diazotrophs to abiotic factors due to microbial associations or interactions present in the field but not in laboratory studies.

Additional resources were also not accounted for in our model since these effects have been poorly constrained in cultures. Trace elements such as nickel involved in photooxidative stress relief, complex forms of organic and organically bound molecules such dissolved organic phosphorus and siderophores, and host-specific requirements, such as silica for DDAs, have been shown to affect diazotroph growth rates (Basu et al., 2019; Benavides et al., 2017; Dyhrman et al., 2006; Ho, 2013).

**Table S3.** Compilation of the ecophysiological parameters influencing growth rates of cyanobacterial diazotrophs in cultures. All parameters are based on changes in relative growth rates unless otherwise noted.

| Genus strain | Temperature | | Light | | Nutrient limitation | |
|---|---|---|---|---|---|---|
| | $\mu_{max}$ ($d^{-1}$) | $Q_{10}$ ($T_{opt}$) unitless | $E_k$ ($\mu$mol quanta $m^{-2}$ $s^{-1}$) | $E_c$ | $k_{PO_4^{3-}}$ (nmol $L^{-1}$) | $k_{Fe}$ |
| *Crocosphaera* WH0401 (small-size) | 0.58[m] | 1.57 ± 0.19[c] (30 °C) | 84 ± 2.2[m] | 19 ± 7[m] | | |
| WH0003 (large-size) | 0.86[d] | 1.33 ± 0.28[c] (30 °C) | 78 ± 4.4[d] | 22 ± 4[d] | 75 ± 12.4[i] | 0.35[j] (WH8501) |
| *Richelia* | 0.60[f] | | 30 ± 1.7[f] | 9 ± 2[f] | 130[l] | 0.5[l] |
| *Calothrix* | 0.26[g] | | 232 ± 5[h] | -182 ± 42[h**] | 130[l] | 0.5[l] |
| *Trichodesmium* IMS101 | 0.33[e] | 2.31 ± 0.07[n] (28°C) | 149 ± 33[e] | 14 ± 9[e] | 50 ± 24[i] | 0.14[k] |

[a]: Brauer et al. (2013) [b]: Taniuchi et al. (2012) [c]: Fu et al. (2014) [d]:N. S. Garcia et al. (2013) [e]: Boatman et al. (2017) [f]: Villareal (1990) Note that $\mu_{max}$ ($d^{-1}$) was derived from division rates of 0.87 div $d^{-1}$x $\log_e(2)$ [g]: Foster et al. (2010) [h**]: Foster et al. (2010) Note that parameters are based on $N_2$ fixation rates not growth rates [i]: Garcia et al. (2015) for cultures maintained at low dFe concentration (0.12-0.35 nM) [j]: Jacq et al. (2014) for a strain of ~3.5 $\mu$m according to Webb et al. (2009) [k]: Boatman et al. (2018) [l]: Follett et al. (2018) [m]: N. Garcia et al. (2013) [n]: Breitbarth et al. (2007). Definition of the parameters: $\mu_{max}$ (theoretical maximum growth rate, Eq. 1), $Q_{10}$ (Arrhenius coefficient, Eq. 2), $T_{opt}$ (optimal temperature, Eq. 2), $E_k$ (light saturation coefficient, Eq. 4), $E_c$ (light compensation coefficient, Eq. 3), $k_{PO_4^{3-}}$ / $k_{Fe}$ (half-saturation coefficients, Eq. 5). Note that $k_{Fe}$ is based on culture kinetics of dissolved inorganic iron (Fe') uptake for *Crocosphaera* and *Trichodesmium*, which is assumed to be the only form of iron bioavailable, and from allometric scaling for DDAs.

**Text S3: Sensitivity analysis of the predator-prey model**

We performed a sensitivity analysis on the predator-prey model coefficients in order to determine the optimal timestep of the multi-linear regression. Numerous nano- and micro-plankton, including the diazotrophs *Crocosphaera, Trichodesmium*, and DDAs in non-bloom periods, as well as their protistan grazers, can be present at low abundances in the North Pacific Gyre (Pasulka et al., 2013; Venrick, 1974). Since the model is based on the dynamics of both diazotrophs and putative grazers as detected by the IFCb, it is important to account for their low abundances and optimize the accuracy of the IFCb observations. One way to increase this accuracy is by binning consecutive samples in order to increase the volume analyzed (5 mL per single sample) and subsequently decrease the detection limit (corresponding to 200 cells $L^{-1}$ in single sample). A number of putative grazers identified by the predator-prey model had abundances <200 cells $L^{-1}$ (see Fig. 7 in the main text); hence, we needed to decrease the detection limit accordingly. As this is equivalent to decreasing the temporal resolution of the timeseries used to fit the model, we anticipated a trade-off between the increase of the timestep (and accuracy of the timeseries used in the model) and the decrease of the number of observations.

A low number of observations may be particularly problematic when fitting a multi-linear model with a large number of variables (in our case 103 and 89 populations in 2017 and 2018); hence, we estimated the uncertainty, expressed as a percentage of the predator-prey interaction coefficients fitted by the linear model, as a function of the model timestep. We propagated the uncertainties of the individual coefficients to estimate the overall uncertainty of grazing rates on specific diazotrophs (calculated as the sum of individual grazing coefficients). Results are shown in Fig. S1 for timesteps of 20 mins (the highest temporal resolution of the IFCb), 40 mins, 1, 2, and 4 hours. Beyond 4 hours, most coefficients become non-significant due to the imbalance between the number of observations and the number of variables. In general, uncertainties for cell-specific grazing rates ranged between 3 and 30%, with the exception of copepods, which had the lowest abundance (1.6-9.9 cells $L^{-1}$). As a result, the overall uncertainty of grazing rate estimates of *Trichodesmium* exceeded 50%, and we discarded these estimates to calculate the overall grazing rates reported in Table 2. As expected, the overall uncertainty was minimal, 12 ± 6% on average, after binning 2 or 3 consecutive samples (temporal resolution of 1 hour) for most interactions, so we decided to fit the model at that temporal resolution.

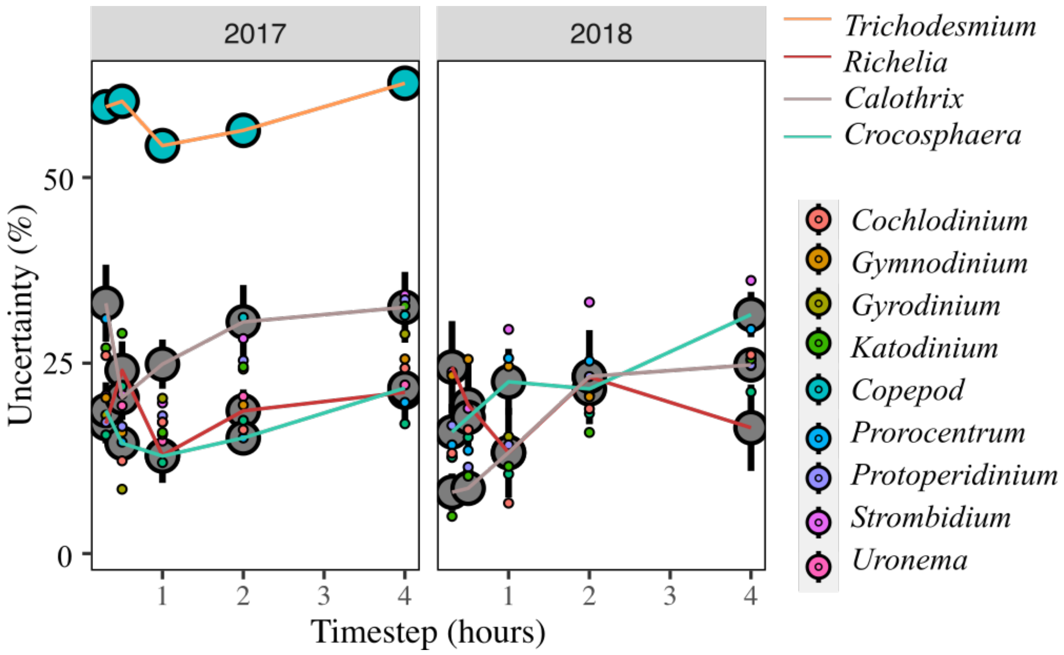


**Figure S1:** Sensitivity analysis of the uncertainty in grazing rate estimates fitted by the predator-prey model. The optimal timestep was determined based on the minimal uncertainty, expressed as a percentage of the linear interaction coefficients, of individual grazing coefficients. The figure highlights the trade-offs between a high temporal resolution (20 mins), yielding to a high number of observations but inaccurate abundances for rare taxa, and coarse resolution (4 hours), yielding more accurate abundance estimates for most taxa but a low number of observations.

**Text S4: Analysis of the IFCb fluorescence of *Trichodesmium* filaments across eddies**

We analyzed the properties of *Trichodesmium* filaments as a proxy for senescence in the 2017 eddies, as well as in bloom samples collected at the frontal station in 2017. The number of observations was too low in 2018 to perform this analysis. Both cell size shrinkage and loss of pigments have been reported as potential markers for programmed cell death (PCD) of viral lysis (Berman-Frank et al., 2004; Hewson et al., 2004). In the absence of direct markers, such as specific viral loads or PCD assays, we used the size and red fluorescence measured by the IFCb to assess evidence for scenescent *Trichodesmium* populations (Fig. S2A). Red fluorescence, emitted by chlorophyll and phycocyanin pigments, generally decreases during daylight due to non-photochemical quenching (data not shown); hence, we only used the surface samples collected at nighttime within eddy centers. Luckily, we encountered the *Trichodesmium* bloom at the frontal station near dusk and proceeded to analyze the bucket samples within 2 hours (first sample analyzed at 18:11 local time). The average red fluorescence measured by the IFCb scales to the entire filament, thus we normalized the average fluorescence by the filament biovolume, in order to account for the various size of trichomes (Fig. S2B). The average normalized fluorescence did not differ between eddy centers, but significantly decreased in the bloom samples (Pearson's chi-squared test). The log-transformed fluorescence followed a bimodal distribution in all samples, corresponding to dimly- and highly-fluorescing *Trichodesmium* filaments (Fig. S2C). We estimated the percentage of dimly-fluorescing *Trichodesmium*, as a proxy for senescent filaments, using a 2-components mixture Gaussian model (R function ‘mclust’, G=2). A majority (65%) of the trichomes were dimly-fluorescing in the bloom sample, compared to 15% and 18% in the anticyclone and cyclone respectively.

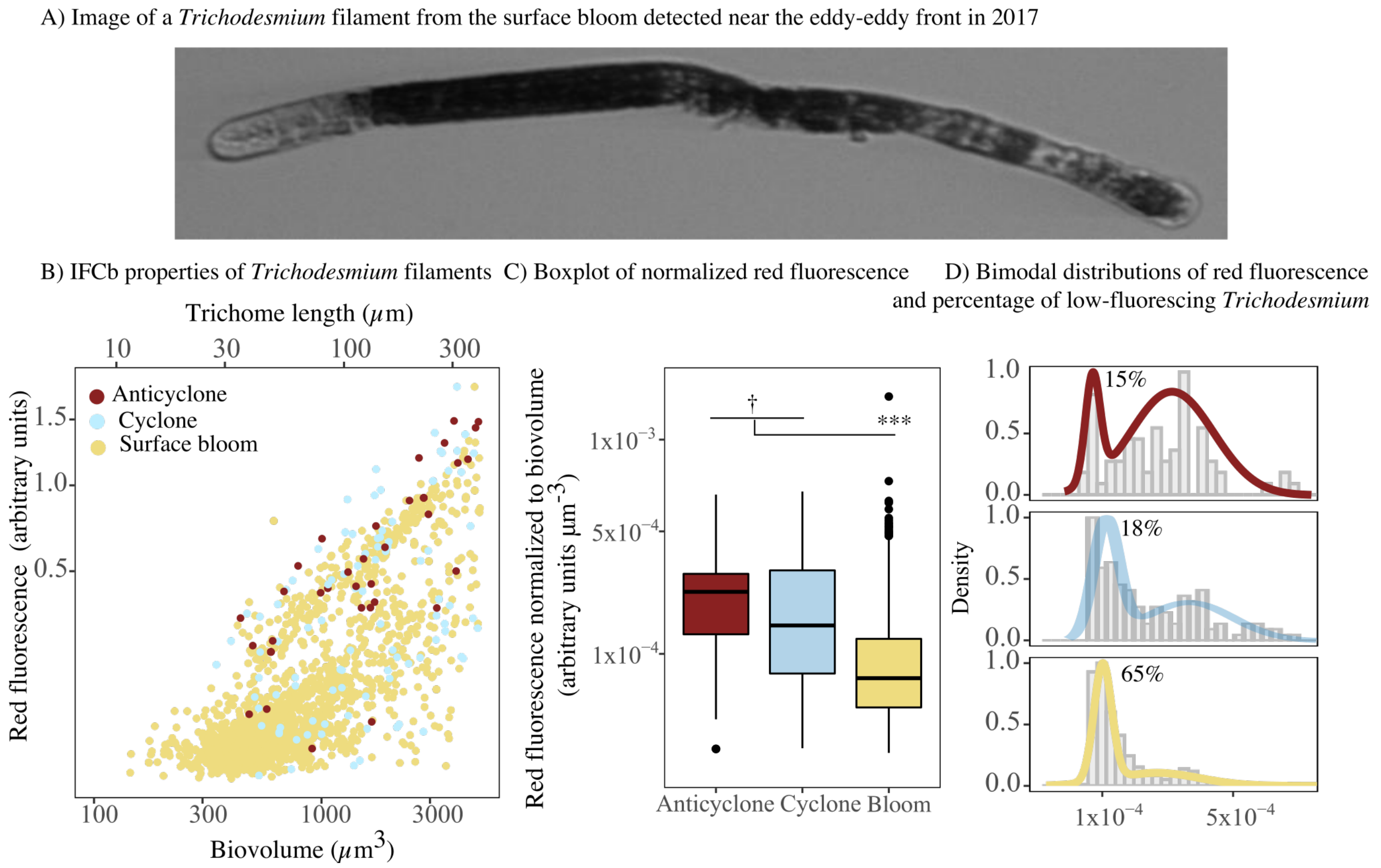


**Figure S2:** Properties of *Trichodesmium* filaments measured by the IFCb in the eddy centers and in surface bloom samples in 2017. Images (A) and filament-specific scatterplot of *Trichodesmium* size against red fluorescence (B) showed that numerous filaments were smaller and lacked pigments in the surface bloom. Normalized red fluorescence did not differ between eddy centers, but was significantly lower in surface bloom (C). The log-transformed fluorescence followed a bimodal distribution, distinguishing the dimly- and highly-fluorescing trichomes (D).

**Text S5: Distributions of diazotroph putative grazers in eddy cruises**

In this section, we present the patterns of grazing rate estimates, derived from the sum of the product of individual interactions and abundances of specific diazotrophs, along with the relative abundances of expressed sequence reads (ribotags) of the putative grazers of diazotrophs identified by the predator-prey model and IFCb images. Ribotags from near the V4 region of the 18S rRNA sequences in 2017 metatrancriptome libraries were analyzed, with raw sequence reads available on NCBI SRA under project number PRJNA515070. Since grazing rate estimates vary as a function of both grazer abundance and cell-specific activity, we leveraged metatranscriptomes (which also vary with abundance and/or activity) from the 2017 cruise to compare the patterns observed in eddies of specific polarity.

On the 2017 cruise, eukaryotic metatranscriptomic samples (20 L) were collected in triplicate every day at ~14:00 local time within the mixed layer (15 m depth). Seawater was subsampled from the CTD rosette and prescreened through a 200 $\mu$m Nylon® mesh, then filtered onto two 5 $\mu$m polycarbonate filters (47 mm) using a peristaltic pump. Total RNA was extracted from individual filters following methods described in Harke et al. (2019). Briefly, the extracted total RNA was sequenced directly (herein referred to as “unselected reads”) via Illumina HiSeq 2000 sequencing, targeting 60 million paired-end reads, at the JP Sulzberger Columbia Genome Center.

Unselected 18S rRNA sequence reads were analyzed using RiboTagger (Xie et al., 2016) which uses a position-specific scoring matrix to scan all rRNA reads to detect sequences next to the V4 region of 18S rRNA (Silva version 119; Quast et al., 2013), herein called ‘ribotags’. Ribotags matching the identity of putative diazotrophic grazers were extracted at the genus level, or at the next highest taxonomic identification level if not possible (e.g. copepod) after Harke et al. (2021). A few putative grazers were not identified in the ribotag dataset (e.g. *Uronema* and *Katodinium*).

The grazing rate estimates and ribotag data were grouped by hierarchical cluster analysis (R function ‘hclust’), using the Euclidian dissimilarities between grazing rate estimates (R function ‘dist’), to identify which taxa consistently displayed a relative increase, relative decrease, or inconsistent patterns in cyclones and anticyclones. We tested the differences in grazing rates estimates (*g*) and ribotag concentrations between eddy types by generalized linear regression, assuming a quasi-Binomial and quasi-Poisson distributions for the rates and concentrations respectively, weighted by the uncertainties of the estimates with the R function 'glm'.

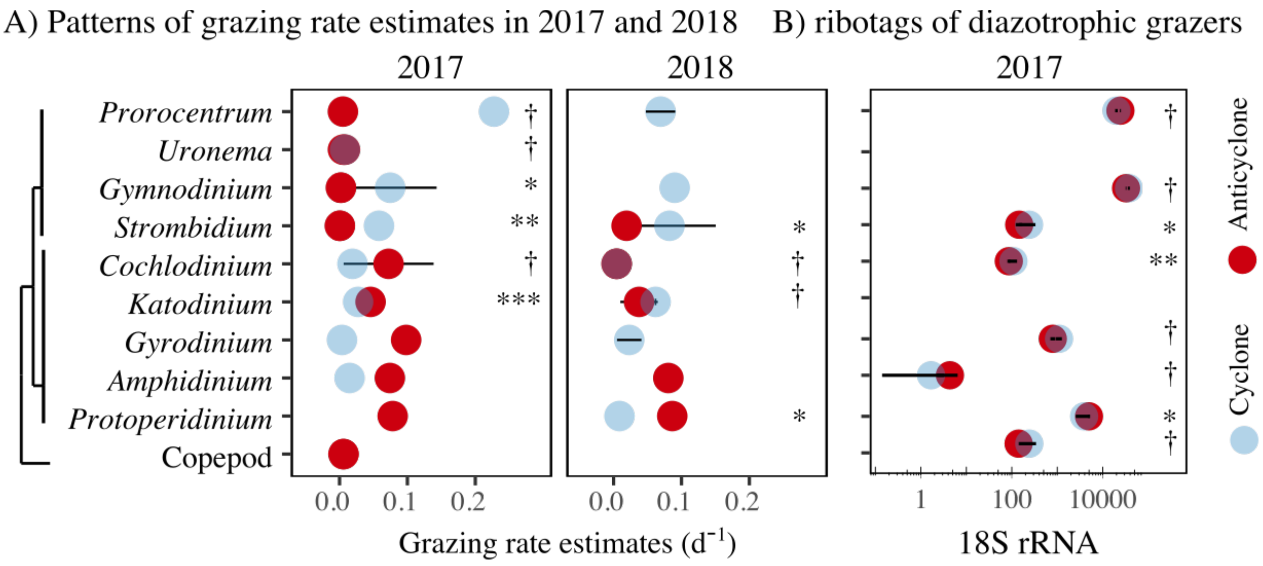


**Figure S3:** Patterns of grazing rate estimates fitted by a predator-prey model in 2017 and 2018 (A) and ribotags of putative diazotrophic grazers in 2017 (B). Protistan grazers were sorted by hierarchical clustering to identify the taxa presenting elevated grazing or ribotag in cyclones (i.e. *Gymnodinium, Strombidium*), or in anticyclones (i.e. *Katodinium, Amphidinium, Protoperidinium*). Differences were tested by a generalized linear model and highlighted some taxa with inconsistent patterns between eddies, such as *Cochlodinium* and copepods. *P*-values: *** (<0.001) ** (<0.005) * (<0.01) † (not significant)

**Table S4:** *nifH* gene abundance data from all sampled stations in this study. Values for the 2018 cruise represent the average ± standard deviation from two biological replicates; biological replicates were not collected during the 2017 cruise. All measured values are reported, including those below detection limits. Detection limits, calculated as 3 times the standard deviation of no template control reactions, were evaluated separately for each assay as follows: UCYN-A1, 1.3; UCYN-B, 3.5; UCYN-C, 0.9; *Trichodesmium*, 3.3; Het-1, 3.6; Het-2, 3.8; and Het-3, 0.9 *nifH* gene copies per µL DNA in each ddPCR reaction. Mean volumetric detection limits for each assay are displayed on Fig. 3.

*See "Supporting Information Tables S2 and S4" excel spreadsheet*

### References

Barone, B., Church, M. J., Dugenne, M., Hawco, N. J., Jahn, O., White, A. E., John, S. G., Follows, M. J., DeLong, E. F., & Karl, D. M. (2022). Biogeochemical dynamics in adjacent mesoscale eddies of opposite polarity. *Global Biogeochemical Cycles, 36*(2), e2021GB007115.

Basu, S., Gledhill, M., de Beer, D., Matondkar, S. P., & Shaked, Y. (2019). Colonies of marine cyanobacteria *Trichodesmium* interact with associated bacteria to acquire iron from dust. *Communications biology, 2*(1), 1-8.

Benavides, M., Berthelot, H., Duhamel, S., Raimbault, P., & Bonnet, S. (2017). Dissolved organic matter uptake by *Trichodesmium* in the Southwest Pacific. *Scientific Reports, 7*(1), 1-6.

Bench, S. R., Heller, P., Frank, I., Arciniega, M., Shilova, I. N., & Zehr, J. P. (2013). Whole genome comparison of six *Crocosphaera watsonii* strains with differing phenotypes. *Journal of Phycology, 49*(4), 786-801.

Berman-Frank, I., Bidle, K. D., Haramaty, L., & Falkowski, P. G. (2004). The demise of the marine cyanobacterium, *Trichodesmium* spp., via an autocatalyzed cell death pathway. *Limnology and Oceanography, 49*(4), 997-1005.

Boatman, T. G., Lawson, T., & Geider, R. J. (2017). A key marine diazotroph in a changing ocean: The interacting effects of temperature, $CO_2$ and light on the growth of *Trichodesmium erythraeum* IMS101. *PloS One, 12*(1), e0168796.

Boatman, T. G., Oxborough, K., Gledhill, M., Lawson, T., & Geider, R. J. (2018). An integrated response of *Trichodesmium erythraeum* IMS101 growth and photo-physiology to iron, $CO_2$, and light intensity. *Frontiers in Microbiology, 9*, 624.

Boyd, P. W., Rynearson, T. A., Armstrong, E. A., Fu, F., Hayashi, K., Hu, Z., Hutchins, D. A., Kudela, R. M., Litchman, E., & Mulholland, M. R. (2013). Marine phytoplankton temperature versus growth responses from polar to tropical waters–outcome of a scientific community-wide study. *PloS One, 8*(5), e63091.

Brauer, V. S., Stomp, M., Rosso, C., van Beusekom, S. A., Emmerich, B., Stal, L. J., & Huisman, J. (2013). Low temperature delays timing and enhances the cost of nitrogen fixation in the unicellular cyanobacterium *Cyanothece*. *The ISME journal, 7*(11), 2105-2115.

Breitbarth, E., Oschlies, A., & LaRoche, J. (2007). Physiological constraints on the global distribution of *Trichodesmium*? Effect of temperature on diazotrophy. *Biogeosciences, 4*(1), 53-61.

Breitbarth, E., Wohlers, J., Kläs, J., LaRoche, J., & Peeken, I. (2008). Nitrogen fixation and growth rates of *Trichodesmium* IMS-101 as a function of light intensity. *Marine Ecology Progress Series, 359*, 25-36.

Bundy, R. M., Boiteau, R. M., McLean, C., Turk-Kubo, K. A., McIlvin, M. R., Saito, M. A., Van Mooy, B. A., & Repeta, D. J. (2018). Distinct siderophores contribute to iron cycling in the mesopelagic at station ALOHA. *Frontiers in Marine Science, 5*, 61.

Caputo, A., Stenegren, M., Pernice, M. C., & Foster, R. A. (2018). A short comparison of two marine planktonic diazotrophic symbioses highlights an un-quantified disparity. *Frontiers in Marine Science, 5*, 2.

de Boyer Montégut, C., Madec, G., Fischer, A. S., Lazar, A., & Iudicone, D. (2004). Mixed layer depth over the global ocean: An examination of profile data and a profile-based climatology. *Journal of Geophysical Research: Oceans, 109*, C12003.

Delmont, T. O. (2021). Discovery of nondiazotrophic *Trichodesmium* species abundant and widespread in the open ocean. *Proceedings of the National Academy of Sciences USA, 118*(46), e2112355118.

Dugenne, M., Henderikx Freitas, F., Wilson, S. T., Karl, D. M., & White, A. E. (2020). Life and death of *Crocosphaera* sp. in the Pacific Ocean: Fine scale predator–prey dynamics. *Limnology and Oceanography, 65*, 2603-2617.

Dyhrman, S., Chappell, P., Haley, S., Moffett, J., Orchard, E., Waterbury, J., & Webb, E. (2006). Phosphonate utilization by the globally important marine diazotroph *Trichodesmium*. *Nature, 439*(7072), 68-71.

Edwards, K. F., Thomas, M. K., Klausmeier, C. A., & Litchman, E. (2015). Light and growth in marine phytoplankton: allometric, taxonomic, and environmental variation. *Limnology and Oceanography, 60*(2), 540-552.

Eppley, R. W. (1972). Temperature and phytoplankton growth in the sea. *Fishery Bulletin, 70*(4), 1063-1085.

Follett, C. L., Dutkiewicz, S., Karl, D. M., Inomura, K., & Follows, M. J. (2018). Seasonal resource conditions favor a summertime increase in North Pacific diatom–diazotroph associations. *The ISME journal, 12*(6), 1543-1557.

Foster, R. A., Goebel, N. L., & Zehr, J. P. (2010). Isolation of *Calothrix rhizosoleniae* (cyanobacteria) strain SC01 from *Chaetoceros* (Bacillariophyta) spp. diatoms of the subropical North Pacific Ocean. *Journal of Phycology, 46*(5), 1028-1037.

Foster, R. A., Kuypers, M. M., Vagner, T., Paerl, R. W., Musat, N., & Zehr, J. P. (2011). Nitrogen fixation and transfer in open ocean diatom–cyanobacterial symbioses. *The ISME journal, 5*(9), 1484-1493.

Foster, R. A., Sztejrenszus, S., & Kuypers, M. M. (2013). Measuring carbon and $N_2$ fixation in field populations of colonial and free-living unicellular cyanobacteria using nanometer-scale secondary ion mass spectrometry. *Journal of Phycology, 49*, 502-516.

Frischkorn, K. R., Haley, S. T., & Dyhrman, S. T. (2018). Coordinated gene expression between *Trichodesmium* and its microbiome over day–night cycles in the North Pacific Subtropical Gyre. *The ISME journal, 12*(4), 997-1007.

Frischkorn, K. R., Rouco, M., Van Mooy, B. A., & Dyhrman, S. T. (2017). Epibionts dominate metabolic functional potential of *Trichodesmium* colonies from the oligotrophic ocean. *The ISME journal, 11*(9), 2090-2101.

Fu, F.-X., Yu, E., Garcia, N. S., Gale, J., Luo, Y., Webb, E. A., & Hutchins, D. A. (2014). Differing responses of marine $N_2$ fixers to warming and consequences for future diazotroph community structure. *Aquatic Microbial Ecology, 72*, 33-46.

Fu, F. X., Zhang, Y., Bell, P. R., & Hutchins, D. A. (2005). Phosphate uptake and growth kinetics of *Trichodesmium* (Cyanobacteria) isolates from the North Atlantic Ocean and The Great Barrier Reef, Australia. *Journal of Phycology, 41*(1), 62-73.

Garcia, N., Fu, F., Breene, C., Elizabeth, K., Bernhardt, P., Mulholland, M., & Hutchins, D. (2013). Combined effects of $CO_2$ and light on large and small isolates of the unicellular $N_2$-fixing cyanobacterium *Crocosphaera watsonii* from the western tropical Atlantic Ocean. *European Journal of Phycology, 48*(1), 128-139.

Garcia, N. S., Fu, F., Sedwick, P. N., & Hutchins, D. A. (2015). Iron deficiency increases growth and nitrogen-fixation rates of phosphorus-deficient marine cyanobacteria. *The ISME journal, 9*(1), 238-245.

Garcia, N. S., Fu, F.-X., & Hutchins, D. A. (2013). Colimitation of the unicellular photosynthetic diazotroph *Crocosphaera watsonii* by phosphorus, light, and carbon dioxide. *Limnology and Oceanography, 58*(4), 1501-1512.

Gradoville, M. R., Crump, B. C., Church, M. J., Letelier, R. M., & White, A. E. (2017). Microbiome of *Trichodesmium* colonies from the North Pacific Subtropical Gyre. *Frontiers in Microbiology, 8*, 1122.

Guo, M., Xiu, P., Chai, F., & Xue, H. (2019). Mesoscale and submesoscale contributions to high sea surface chlorophyll in subtropical gyres. *Geophysical Research Letters, 46*(22), 13217-13226.

Harke, M. J., Frischkorn, K. R., Haley, S. T., Aylward, F. O., Zehr, J. P., & Dyhrman, S. T. (2019). Periodic and coordinated gene expression between a diazotroph and its diatom host. *The ISME journal, 13*(1), 118-131.

Harke, M. J., Frischkorn, K. R., Hennon, G. M., Haley, S. T., Barone, B., Karl, D. M., & Dyhrman, S. T. (2021). Microbial community transcriptional patterns vary in response to mesoscale forcing in the North Pacific Subtropical Gyre. *Environmental Microbiology, 23*(8), 4807-4822.

Hewson, I., Govil, S. R., Capone, D. G., Carpenter, E. J., & Fuhrman, J. A. (2004). Evidence of *Trichodesmium* viral lysis and potential significance for biogeochemical cycling in the oligotrophic ocean. *Aquatic Microbial Ecology, 36*(1), 1-8.

Hewson, I., Poretsky, R. S., Dyhrman, S. T., Zielinski, B., White, A. E., Tripp, H. J., Montoya, J. P., & Zehr, J. P. (2009). Microbial community gene expression within colonies of the diazotroph, *Trichodesmium*, from the Southwest Pacific Ocean. *The ISME journal, 3*(11), 1286-1300.

Hmelo, L. R., Van Mooy, B. A. S., & Mincer, T. J. (2012). Characterization of bacterial epibionts on the cyanobacterium Trichodesmium. *Aquatic Microbial Ecology, 67*(1), 1-14.

Ho, T.-Y. (2013). Nickel limitation of nitrogen fixation in *Trichodesmium*. *Limnology and Oceanography, 58*(1), 112-120.

Hutchins, D. A., Fu, F.-X., Webb, E. A., Walworth, N., & Tagliabue, A. (2013). Taxon-specific response of marine nitrogen fixers to elevated carbon dioxide concentrations. *Nature geoscience, 6*(9), 790-795.

Hynes, A. M. (2009). *Diversity of the marine Cyanobacterium Trichodesmium: characterization of the Woods Hole culture collection and quantification of field populations* [Doctoral dissertation, Massachusetts Institute of Technology].

Hynes, A. M., Webb, E. A., Doney, S. C., & Waterbury, J. B. (2012). Comparison of cultured *Trichodesmium* (Cyanophyceae) with species characterized from the field. *Journal of Phycology, 48*, 196-210.

Jacq, V., Ridame, C., l'Helguen, S., Kaczmar, F., & Saliot, A. (2014). Response of the unicellular diazotrophic cyanobacterium *Crocosphaera watsonii* to iron limitation. *PloS One, 9*(1), e86749.

Litchman, E., Edwards, K. F., Klausmeier, C. A., & Thomas, M. K. (2012). Phytoplankton niches, traits and eco-evolutionary responses to global environmental change. *Marine Ecology Progress Series, 470*, 235-248.

McGillicuddy Jr, D. J. (2014). Do *Trichodesmium* spp. populations in the North Atlantic export most of the nitrogen they fix? *Global Biogeochemical Cycles, 28*(2), 103-114.

Momper, L. M., Reese, B. K., Carvalho, G., Lee, P., & Webb, E. A. (2015). A novel cohabitation between two diazotrophic cyanobacteria in the oligotrophic ocean. *The ISME journal, 9*, 882-893.

Montoya, J. P., Voss, M., Kahler, P., & Capone, D. G. (1996). A simple, high-precision, high-sensitivity tracer assay for $N_2$ fixation. *Applied and Environmental Microbiology, 62*(3), 986-993.

Orchard, E., Webb, E., & Dyhrman, S. (2003). Characterization of phosphorus-regulated genes in *Trichodesmium* spp. *The Biological Bulletin, 205*(2), 230-231.

Pasulka, A. L., Landry, M. R., Taniguchi, D. A., Taylor, A. G., & Church, M. J. (2013). Temporal dynamics of phytoplankton and heterotrophic protists at station ALOHA. *Deep Sea Research Part II: Topical Studies in Oceanography, 93*, 44-57.

Pyle, A. E., Johnson, A. M., & Villareal, T. A. (2020). Isolation, growth, and nitrogen fixation rates of the Hemiaulus-Richelia (diatom-cyanobacterium) symbiosis in culture. *PeerJ, 8*, e10115.

Qu, P., Fu, F.-X., Kling, J. D., Huh, M., Wang, X., & Hutchins, D. A. (2019). Distinct responses of the nitrogen-fixing marine cyanobacterium *Trichodesmium* to a thermally variable environment as a function of phosphorus availability. *Frontiers in Microbiology, 10*, 1282.

Quast, C., Pruesse, E., Yilmaz, P., Gerken, J., Schweer, T., Yarza, P., Peplies, J., & Glöckner, F. O. (2013). The SILVA ribosomal RNA gene database project: improved data processing and web-based tools. *Nucleic Acids Research, 41*(D1), D590-D596.

Rouco, M., Frischkorn, K. R., Haley, S. T., Alexander, H., & Dyhrman, S. T. (2018). Transcriptional patterns identify resource controls on the diazotroph *Trichodesmium* in the Atlantic and Pacific oceans. *The ISME journal, 12*(6), 1486-1495.

Rouco, M., Haley, S. T., Alexander, H., Wilson, S. T., Karl, D. M., & Dyhrman, S. T. (2016). Variable depth distribution of *Trichodesmium* clades in the North Pacific Ocean. *Environmental Microbiology Reports, 8*(6), 1058-1066.

Rouco, M., Haley, S. T., & Dyhrman, S. T. (2016). Microbial diversity within the *Trichodesmium* holobiont. *Environmental Microbiology, 18*(12), 5151-5160.

Saito, M. A., Bertrand, E. M., Dutkiewicz, S., Bulygin, V. V., Moran, D. M., Monteiro, F. M., Follows, M. J., Valois, F. W., & Waterbury, J. B. (2011). Iron conservation by reduction of metalloenzyme inventories in the marine diazotroph *Crocosphaera watsonii*. *Proceedings of the National Academy of Sciences USA, 108*(6), 2184-2189.

Sohm, J. A., Hilton, J. A., Noble, A. E., Zehr, J. P., Saito, M. A., & Webb, E. A. (2011). Nitrogen fixation in the South Atlantic Gyre and the Benguela upwelling system. *Geophysical Research Letters, 38*(16), L16608.

Stukel, M. R., Coles, V. J., Brooks, M., & Hood, R. R. (2014). Top-down, bottom-up and physical controls on diatom-diazotroph assemblage growth in the Amazon River plume. *Biogeosciences, 11*(12), 3259-3278.

Tang, W., Cerdán-García, E., Berthelot, H., Polyviou, D., Wang, S., Baylay, A., Whitby, H., Planquette, H., Mowlem, M., & Robidart, J. (2020). New insights into the distributions of nitrogen fixation and diazotrophs revealed by high-resolution sensing and sampling methods. *The ISME journal*, 1-13.

Taniuchi, Y., Chen, Y. l. L., Chen, H. Y., Tsai, M. L., & Ohki, K. (2012). Isolation and characterization of the unicellular diazotrophic cyanobacterium Group C TW3 from the tropical western Pacific Ocean. *Environmental Microbiology, 14*(3), 641-654.

Tian, R., & Chen, C. (2006). Influence of model geometrical fitting and turbulence parameterization on phytoplankton simulation in the Gulf of Maine. *Deep Sea Research Part II: Topical Studies in Oceanography, 53*(23-24), 2808-2832.

Tuo, S.-H., Lee Chen, Y.-L., Chen, H.-Y., & Chen, T.-Y. (2017). Free-living heterocystous cyanobacteria in the tropical marginal seas of the western North Pacific. *Journal of Plankton Research, 39*(3), 404-422.

Venrick, E. (1974). The distribution and significance of *Richelia intracellularis* Schmidt in the North Pacific Central Gyre *Limnology and Oceanography, 19*(3), 437-445.

Villareal, T., & Carpenter, E. (2003). Buoyancy regulation and the potential for vertical migration in the oceanic cyanobacterium *Trichodesmium*. *Microbial Ecology, 45*(1), 1-10.

Villareal, T. A. (1989). Division cycles in the nitrogen-fixing *Rhizosolenia* (Bacillariophyceae)-*Richelia* (Nostocaceae) symbiosis. *British Phycological Journal, 24*(4), 357-365.

Villareal, T. A. (1990). Laboratory culture and preliminary characterization of the nitrogen-fixing *Rhizosolenia-Richelia* symbiosis. *Marine Ecology, 11*(2), 117-132.

Villareal, T. A., Brown, C. G., Brzezinski, M. A., Krause, J. W., & Wilson, C. (2012). Summer diatom blooms in the North Pacific subtropical gyre: 2008–2009. *PloS One, 7*(4), e33109.

Villareal, T. A., & Carpenter, E. J. (1989). Nitrogen fixation, suspension characteristics, and chemical composition of *Rhizosolenia* mats in the central North Pacific gyre. *Biological Oceanography, 6*(3-4), 327-345.

Walworth, N. G., Fu, F.-X., Webb, E. A., Saito, M. A., Moran, D., Mcllvin, M. R., Lee, M. D., & Hutchins, D. A. (2016). Mechanisms of increased *Trichodesmium* fitness under iron and phosphorus co-limitation in the present and future ocean. *Nature communications, 7*(1), 1-11.

Webb, E. A., Ehrenreich, I. M., Brown, S. L., Valois, F. W., & Waterbury, J. B. (2009). Phenotypic and genotypic characterization of multiple strains of the diazotrophic cyanobacterium, *Crocosphaera watsonii*, isolated from the open ocean. *Environmental Microbiology, 11*(2), 338-348.

White, A. E., Spitz, Y. H., Karl, D. M., & Letelier, R. M. (2006). Flexible elemental stoichiometry in *Trichodesmium* spp. and its ecological implications. *Limnol. Oceanogr., 51*, 1777-1790.

Xie, C., Goi, C. L. W., Huson, D. H., Little, P. F., & Williams, R. B. (2016). RiboTagger: fast and unbiased 16S/18S profiling using whole community shotgun metagenomic or metatranscriptome surveys. *BMC Bioinformatics, 17*(19), 277-282.
